# Quantitative trait loci mapped for TCF21 binding, chromatin accessibility and chromosomal looping in coronary artery smooth muscle cells reveal molecular mechanisms of coronary disease loci

**DOI:** 10.1101/2020.02.03.932368

**Authors:** Quanyi Zhao, Michael Dacre, Trieu Nguyen, Milos Pjanic, Boxiang Liu, Dharini Iyer, Paul Cheng, Robert Wirka, Juyong Brian Kim, Hunter B Fraser, Thomas Quertermous

**Author notes:** **Materials and Correspondence** Thomas Quertermous, 300 Pasteur Dr., Falk CVRC, Stanford, CA 94305.

## Abstract

**Background:** To investigate the epigenetic and transcriptional mechanisms of coronary artery disease (CAD) risk, as well as the functional regulation of chromatin structure and function, we have created a catalog of genetic variants associated with three stages of transcriptional *cis*-regulation in primary human coronary artery vascular smooth muscle cells (HCASMC).

**Results:** To this end, we have used a pooling approach with HCASMC lines to map regulatory variation that mediates binding of the CAD associated transcription factor TCF21 with ChIPseq studies (bQTLs), variation that regulates chromatin accessibility with ATACseq studies (caQTLs), and chromosomal looping with HiC methods (clQTLs). We show significant overlap of the QTLs, and their relationship to smooth muscle specific genes and the binding of smooth muscle transcription factors. Further, we use multiple analyses to show that these QTLs are highly associated with CAD GWAS loci and correlated to lead SNPs in these loci where they show allelic effects. We have verified with genome editing that identified functional variants can regulate both chromatin accessibility and chromosomal looping, providing new insights into functional mechanisms regulating chromatin state and chromosomal structure. Finally, we directly link the disease associated *TGFβ1-SMAD3* pathway to the CAD associated *FN1* gene through a response QTL that modulates both chromatin accessibility and chromosomal looping.

**Conclusions:** Together, these studies represent the most thorough mapping of multiple QTL types in a highly disease relevant primary cultured cell type, and provide novel insights into their functional overlap and mechanisms that underlie these genomic features and their relationship to disease risk.

## Background

While there has been considerable success identifying loci in the human genome that are associated with a broad range of human diseases, including coronary artery disease (CAD) [1–3], most identified regions represent non-exonic regulatory sequence and are thus difficult to associate to a particular gene or functional annotation of the genome. Toward this end, numerous studies have explored the genetics of gene expression and mapped expression quantitative trait locus variants to inform on which SNPs and which genes are causally related to disease in the associated loci [4–7]. A number of these expression quantitative trait variants (eQTLs) have been found to be the causal variants in disease loci, and in general promoted causal gene discovery [8].

The regulatory variants that modulate other genomic features have also been characterized. QTLs for histone modification and chromatin accessibility, and more recently chromosomal looping have also been mapped [9–12]. Identification of genomic QTLs has been accelerated by use of a pooling approach that significantly reduces the number of individuals that have to be studied, and promoted the mapping of allelic variation that regulates transcription factor binding [13] and chromatin accessibility in a large number of ethnically diverse individuals [9]. Such efforts have provided important information regarding the regulatory structure of the non-coding sequence in the genome and identified how such variation may regulate the risk for complex human disease. However, in general these studies conducted with large high throughput genomic datasets have not been validated with targeted follow-up studies, and the ramifications of their allele-specific effects not investigated. Also, these studies have primarily been conducted in lymphoblastoid cell lines and restricted to a single type of QTL, and thus have not been examined in the context of overlap of QTL functions and their relevance for a broad range of human diseases.

To better understand the functional basis of regulatory features of the human genome, and to accelerate understanding of the transcriptional mechanisms by which these features contribute to CAD, we have mapped quantitative trait variation in disease relevant primary cultured human coronary artery vascular smooth muscle cells (HCASMC). We have identified QTLs for binding of the CAD-associated transcription factor TCF21 (bQTLs), chromatin accessibility (caQTLs), and chromosomal looping (clQTLs), investigated their relationship to one another, and to eQTLs mapped in the same cell type, as well as their overlap with CAD associated genetic variation.

## Results

### bQTL, caQTL and clQTL calling in pooled HCASMC lines

We obtained primary HCASMC lines from commercial vendors. Genotypes were called with whole genome sequencing or genotyping with Illumina chips, phased and imputed against 1000 Genome phase 3 data before merging [14]. We pooled 65 lines for TCF21 ChIPseq and Hi-C, and 71 lines for ATACseq experiments at passage 4-6 (Fig. 1a, Methods). After sequencing, we obtained more than 400 M reads each for pooled TCF21 ChIPseq and ATACseq libraries and 800 M reads for the pooled Hi-C library. Standard pipelines were employed for each of these genomic approaches. We called 22381 high-confidence binding peaks from TCF21 ChIPseq with fold enrichment >10, *P*-value <10^−20^ cutoffs and 18601 open chromatin regions from ATACseq with fold enrichment >4, *P*-value <10^−10^. Most of these TCF21 binding peaks (19705, >88%) and open chromatin regions (17129, >92%) overlapped with the data previously identified in an individual HCASMC line [15]. For the Hi-C data, in addition to standard data processing, we also assigned sequencing reads to the SNPs genotyped in our pooled HCAMSC lines, generating a total of more than 377 million (M) valid interacted reads pairs. Of these, approximately 21.6 M allele-specific interactions with at least one SNP inside the loop boundaries were identified (Additional file1: Suppl Figs. 1a, 1b). Using these interactions, we were be able to find 7916 loops by FitHiC and 3443 loops by HiCCUPS with a stringent *P*-value cutoff (10^−10^). Here we show an example genome region at 7p15.2, comparing our pooled ATACseq track and Hi-C loops for heterologous cell types with our previously published H3K27ac HiChIP loops in an individual HCASMC line[16], and ENCODE generated ChIA-PET loops (Fig. 1b). Our pooled Hi-C loops showed similar patterns but with a higher resolution and more chromosomal contacts compared to the previously reported data. More importantly, our data showed high resolution (5kb) of allele-specific interactions, revealing the differential chromosomal architectures associated with individual SNPs (Fig. 1c).

**Figure 1.**
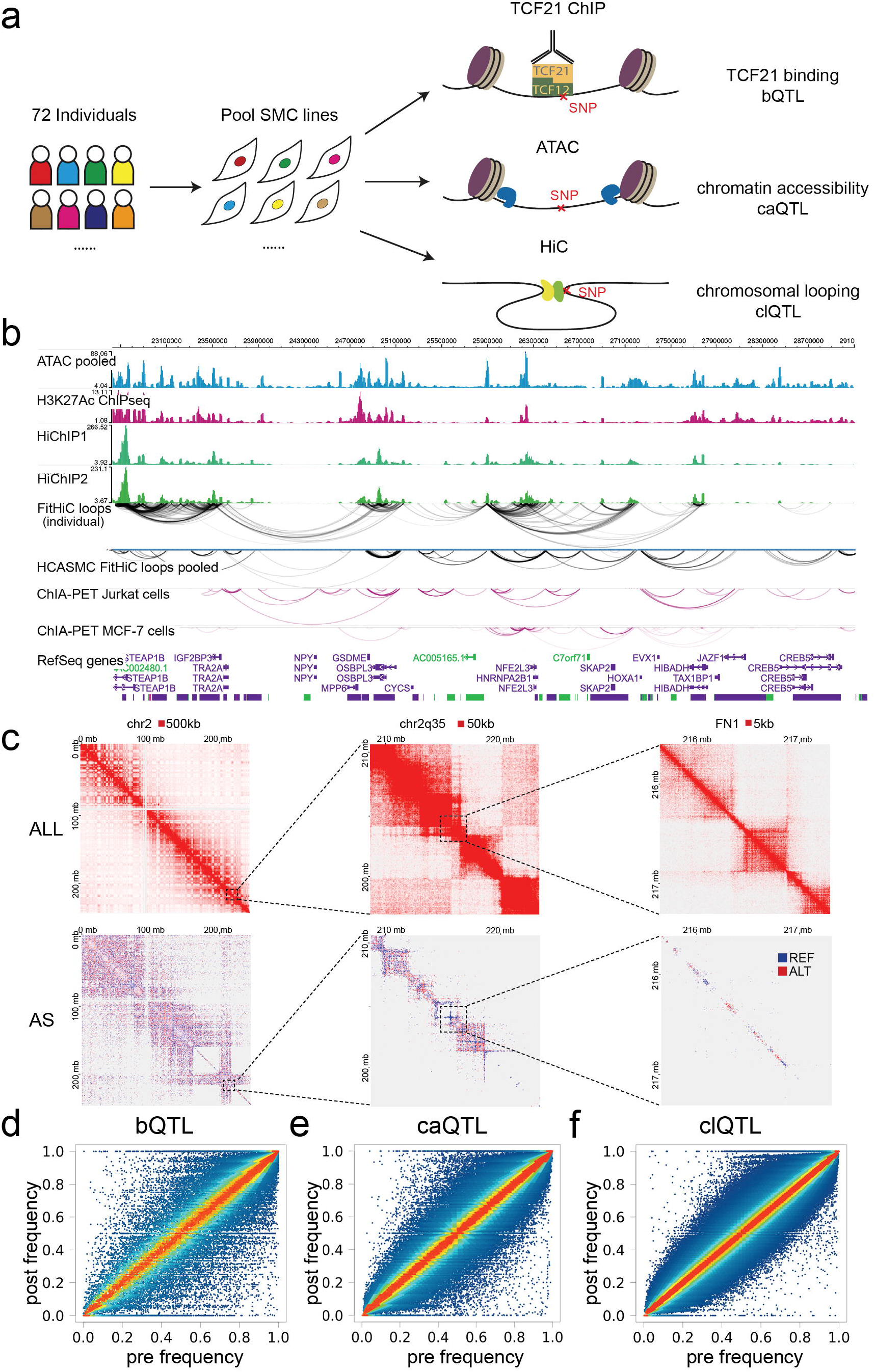
bQTLs, caQTLs and clQTLs in pooled HCASMC lines. (a) Diagram showing the approaches for pooled TCF21 ChIPseq, ATACseq, HiC and QTL calling. (b) Genome browser session showing the HCASMC pooled sequencing tracks, loops called from HiC and their comparison with HiChIP in an individual line or non-SMC ChIA-PET. (c) Heatmaps show the chromosomal contacts on chr2 at a variety of magnifications and resolutions. ALL, all loops; AS, allele-specific loops. (d) Plots showing the regression results and pre-post frequency distributions for TCF21 binding, 5315 bQTLs, (e) chromatin accessibility, 8346 caQTLs and (f) chromosomal looping, 7084 clQTLs,

To call the TCF21 binding (bQTL), chromatin accessibility (caQTL) and chromosomal looping (clQTL) quantitative trait loci, we employed a published regression-based approach which uses post-assay allele frequencies together with genotypes of each sample to infer the proportion of each sample in the pool [9, 13]. Comparing the pre-assay vs. post-assay allele frequencies allows the identification of outlier SNPs where these frequencies are significantly different from one another, which indicates a *cis*-acting QTL variant. Although the pre-post frequency distributions showed a slight reference allele frequency bias after regression, consistent with previously reported studies [13, 16], the majority of variants showed a balanced pattern (Figs. 1d, 1e and 1f). With fold enrichment-optimized [9] at *P* value cutoff 10^−4^, we obtained 5315 significant bQTLs, 8346 caQTLs and 7084 clQTLs with the regression analysis (Additional file 2: Suppl Table. 1).

### QTLs are highly associated with GWAS CAD loci

An initial question we wanted to address was the relationship of mapped QTLs with coronary artery disease (CAD) associated variation. To examine this question, we used GWAS catalog SNPs [17] supplemented with CARDIoGRAMplusC4D variant data from a 1000 Genomes-based meta-analysis [18]. First, we extracted lead SNPs in CAD-associated categories “Atherosclerosis”, “Coronary artery calcification”, “Coronary artery”, “Coronary heart”, “Kawasaki disease” and “Myocardial infarction” from the GWAS catalog along with the lead SNPs in CARDIoGRAMplusC4D. We also randomly selected 27 unrelated diseases to serve as background. We then investigated the enrichment of these lead SNPs in a ±1 kb window around the QTLs.

This analysis revealed that the lead SNPs in CAD-associated categories have strong colocalization (*P*<10^−10^) with the bQTLs, caQTLs and clQTLs. The highest enriched category for bQTL variants was “Myocardial infarction” and for caQTLs “CARDIoGRAMplusC4D” and “Coronary artery calcification”. clQTL variants were enriched for “Myocardial infarction,” “CardiogramplusC4D,” and the highly significant term “Coronary artery,” which was the second most significant category (*P*<10^−100^) ranked by *P*-value (Figs. 2a, 2b and 2c). This finding suggested that these QTLs in HCASMC may play an important role in the genetic mechanism of HCASMC mediated disease progression. All of the QTLs showed enrichment for blood pressure and hypertension phenotypes which are highly consistent with known SMC functions, and association of *TCF21* with blood pressure have been identified by population studies with multiple racial ethnic groups [19–21]. The highly significant enrichment of breast cancer variants in the bQTL category is consistent with TCF21 being a known tumor suppressor and that is dysregulated in breast cancer, but the enrichment of breast cancer variants among the caQTLs and clQTLs is surprising and suggests a similar genomic and genetic architecture between HCASMC and breast cancer risk genes [22].

**Figure 2.**
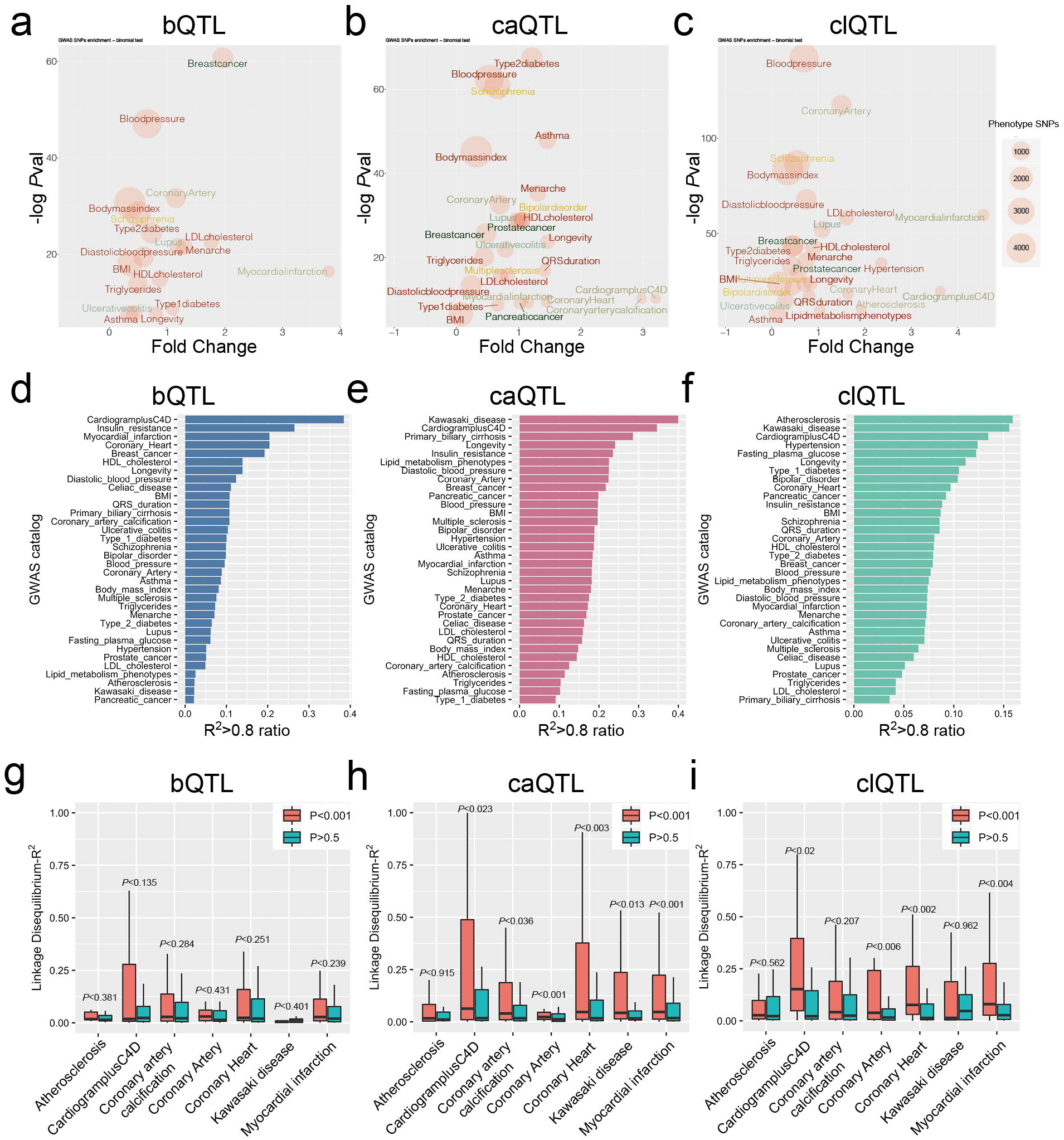
QTLs are highly associated with GWAS CAD loci. (a) bQTL, (b) caQTL and (c) clQTL plots show the lead SNPs of GWAS catalog enrichment in the ±1kb windows of the QTLs, including *P*-values, fold changes and number sizes. (d) bQTL, (e) caQTL and (f) clQTL bar graphs show the ranked GWAS terms for associated lead SNP-QTL pairs with linkage disequilibrium (LD) (R^2^>0.8) in the GWAS catalog, compared to the total number of pairs. (g) bQTL, (h) caQTL and (i) clQTL box plots show the total LD R^2^ distributions of QTL to lead SNP pairs, in GWAS CAD-associated diseases.

To further validate these results, we also evaluated the association of QTLs with disease risk variants by linkage disequilibrium (LD) comparison. We selected all of the GWAS lead SNP-QTL pairs which were separated by less than 100kb, calculated the R^2^ for each pair, and then ranked the categories by the ratio of pairs with R^2^>0.8 compared to the total number of pairs. Each of the three types of QTLs showed CAD related categories as among the three most significant (Figs. 2d, 2e and 2f). Interestingly, these analyses also showed some QTL-specificity: “Myocardial infarction” is enriched in bQTL, “Atherosclerosis” only in clQTL, while “Kawasaki disease” is enriched only for caQTL and clQTL variants (Figs. 2d, 2e and 2f). These results were consistent with the distance-dependent enrichment results (Figs. 2a, 2b and 2c).

We next focused on the CAD-associated categories only, comparing the R^2^ distributions of the LD pairs to the significant QTLs (*P*<10^−4^) with those of non-significant SNPs (*P*>0.5). While the bQTLs showed a trend in the correct direction, they did not show a significant correlation (Fig. 2g). However, the caQTLs (Fig. 2h) and clQTLs (Fig. 2i) showed a statistically greater correlation than non-significant SNPs. The fold change of R^2^>0.8 pairs of significant to non-significant QTLs indicated enrichments for the majority of the cardiovascular diseases (Additional file 1: Suppl Figs. 2a, 2b, and 2c). These data demonstrate that the QTLs we identified are significantly enriched in CAD-associated loci.

### QTLs colocalize with SMC transcription factor binding and epigenetic modifications

We have previously shown that open chromatin regions in HCASMC are enriched for CAD-associated loci and multiple SMC-specific transcription factor (TF) binding sites, such as TCF21 [23]. We extended these studies by investigating TF motif enrichment analysis at the identified QTLs. We created a window of analysis that extended 50 bp upstream and downstream of each significant (*P*<10^−4^) QTL and scanned these sequences by matching to the HOMER known motifs or JASPAR core 2018 vertebrate database [24]. Non-significant QTLs (*P*>0.5) with the same window size served as background. The most enriched motifs around bQTLs were TCF21 and its bHLH Class I dimerization partner TCF12 (Fig. 3a). The AP-1 complex subunit (ATF3 and JUNB), Hippo pathway transcription factor TEAD1 and the chromatin regulator CTCF motifs were also enriched in these regions of the genome, suggesting that they could be regulated together with TCF21 binding at these loci (Fig. 3a). For caQTLs, we found that the AP-1 complex is the dominant TF motif nearby, along with those for SMC functional TFs such as TEAD1/3, SMAD3, TCF21 and ARNT (Fig. 3b). These enrichments and colocalizations extend our previous studies [15, 23]. A similar analysis was also performed with clQTLs, which showed ZEB1 and ZEB2 binding site enrichment and also enrichment for a number of TF motifs that haven’t been reported to be associated with CAD or SMC function (Additional file 1: Suppl Figs. 3a, 3b).

**Figure 3.**
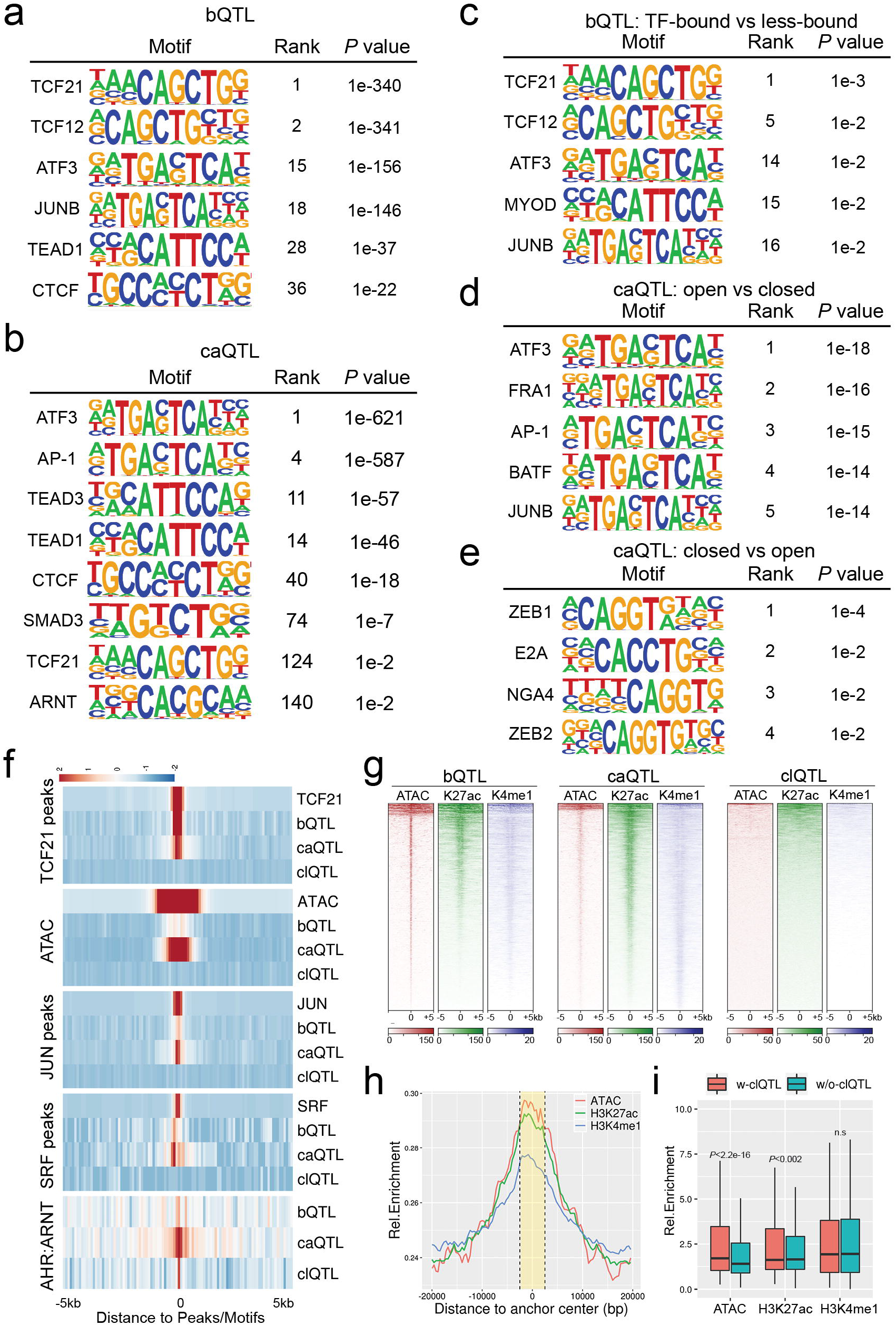
SMC relevant transcription factors are colocalized with QTLs. Results from HOMER reference motif scanning on ±50 bp windows surrounding (a) bQTLs and caQTLs. Allele-specific motif scanning with flipped SNPs in ±20 bp windows surrounding bound-vs-less-bound bQTLs, (d) open-vs-closed caQTLs and (e) closed-vs-open caQTLs. (f) Heatmaps show the bQTL, caQTL and clQTL densities at TCF21 peaks, ATAC peaks, JUN peaks, SRF peaks and AHR:ARNT binding motifs. (g) Heatmaps show the ATACseq open chromatin (ATAC), histone H3K27ac (K27ac) and H3K4me1(K4me1) marks enrichment at bQTLs, caQTLs and clQTLs in ±5 kb windows. (h) Density plot showing open chromatin, H3K27ac and H3K4me1 enrichment at chromosomal loop anchors (yellow area within dashed lines). (i) Boxplots show differences between the enrichments of open chromatin, H3K27ac and H3K4me1, on chromosomal loop anchors with and without clQTLs.

To investigate the allele-specific TF binding at these QTLs, we searched the motifs that were differentially enriched among the high-chromatin accessibility or TCF21 bound alleles compared to the low-chromatin accessibility or TCF21 less bound alleles for the same QTLs in a ±20 bp window. The results show that TCF21/TCF12, AP-1 complex motifs and a bHLH E-box assigned to MYOD are only enriched in open bQTL alleles with no significant TF motifs found in closed alleles (Fig. 3c). The AP-1 complex motif was associated preferentially with open caQTL alleles, presumably through pioneer functions that promote chromatin accessibility (Fig. 3d) [15]. Interestingly, we found accessible ZEB1 and ZEB2 binding sites at closed caQTL alleles, suggesting that they may have the opposite function of AP-1 in SMC (Fig. 3e) [25].

We generated heatmaps to investigate co-localization of the QTLs with each other, and with HCASMC ATACseq and ChIPseq data. TCF21 bQTLs were shown to colocalize with previous ChIPseq data [26] (Fig. 3f) in regions of open chromatin that mediate binding of the SMC related MYOCD-SRF complex [27] and JUN [15], supporting the authenticity of the bQTLs. Additionally, we found ATACseq open chromatin, as well as histone H3K27ac and H3K4me1 marks to be highly enriched at bQTLs (Figs. 3f, 3g), suggesting that they may regulate TCF21 binding by affecting the epigenome and chromatin accessibility. As expected, caQTLs were enriched at ATACseq regions of open chromatin, and further localized to enhancer regions (Figs 3f, 3g). With the exception of AHR:ARNT binding motifs, clQTLs in general showed minimal colocalization with other QTLs or ChIPseq peaks for TFs or histone modifications (Figs. 3f, 3g). However, chromosomal loops were enriched at their anchors for chromatin accessibility as well as H3K27ac and H3K3me1 chromatin marks (Fig. 3h), with those loops having clQTLs in their anchors showing higher chromatin accessibility and H3K27ac levels than those loops without clQTLs (Fig. 3i). Taken together, these data show that variation regulating TCF21 binding colocalizes with caQTLs in regions of open chromatin that have enhancer histone marks, and are enriched for clQTLs at looping anchors, suggesting that epigenetic effects mediate a significant proportion of genomic molecular phenotypes.

### Co-localizing QTLs identify genes that mediate key SMC functional roles

Our next goal was to characterize loci with bQTL, caQTL and clQTL colocalization. We first overlapped these QTLs to examine exact matching and included our published HCASMC eQTL data in this analysis [14]. The number of eQTLs (187299) was comparable to GTEx QTL numbers and QTL-gene pairs (Additional file 1: Suppl Figs. 4a, 4b). We found 446 direct overlaps between bQTLs and caQTLs, 135 between caQTLs and clQTLs, 47 between bQTLs and clQTLs, and 26 overlaps for all three (Fig. 4a). Each QTL group had numerous exact matches with eQTLs while there were few overlaps for all four QTL groups. Interestingly, the overlaps between bQTLs and caQTLs were found to be directional. TF-bound alleles of bQTLs had more matches with open alleles of caQTLs than those with closed alleles while a comparison of less-bound alleles of bQTLs with caQTLs showed opposite results (Additional file 1: Suppl Fig. 4c). We attribute these findings to the colocalization of TCF21 binding and open chromatin regions as we previously described (Figs. 3g, 3h) [15].

**Figure 4.**
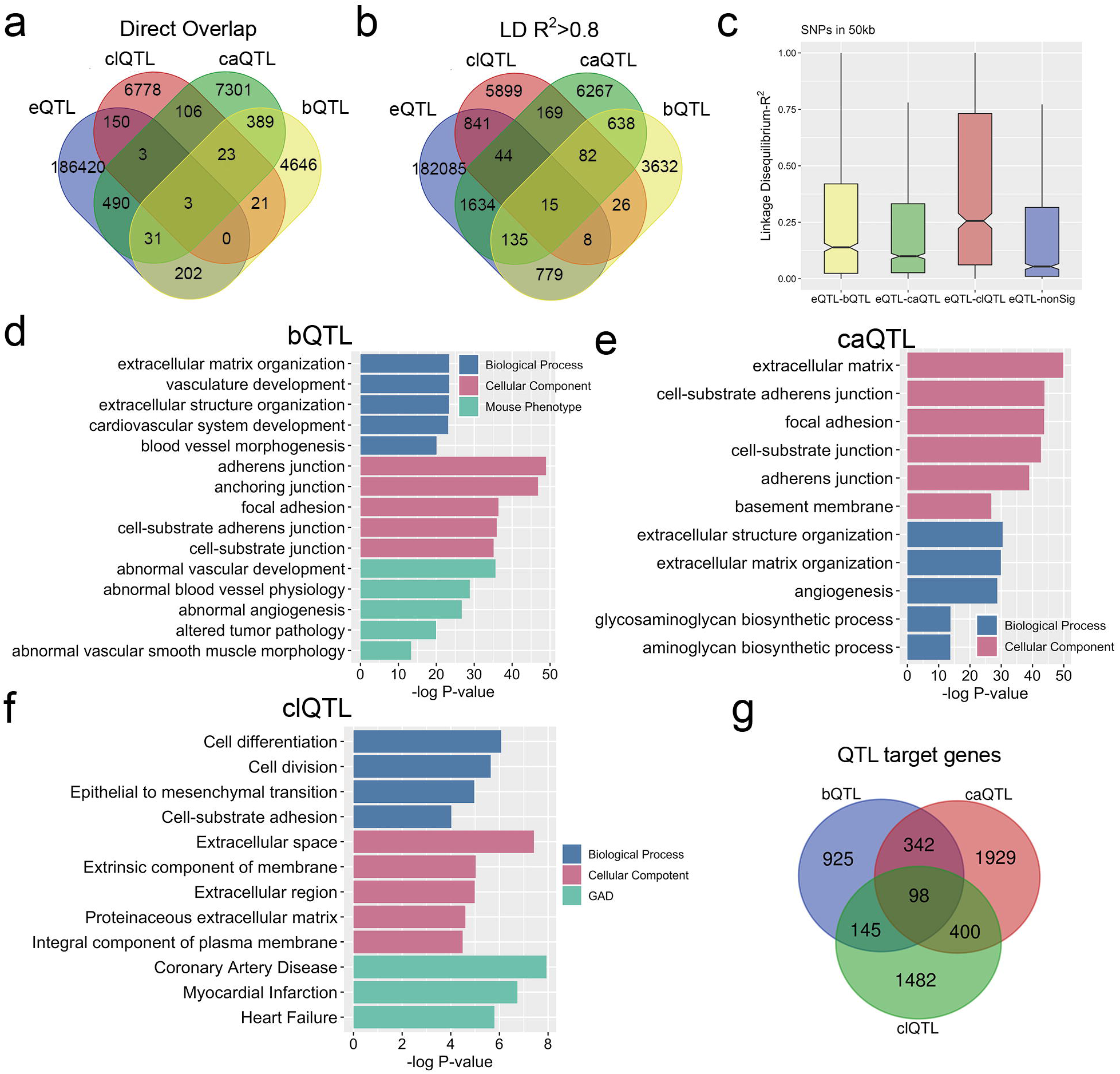
Colocalizing QTLs identify genes that mediate key roles in SMC functions. Venn diagrams show (a) direct overlaps or (b) linkage disequilibrium (LD) R^2^>0.8 between bQTL, caQTL, clQTL and eQTLs all mapped in HCASMC. (c) Box plots show LD R^2^ distributions between bQTL-eQTL, caQTL-eQTL and clQTL-eQTL pairs. Bar graphs show Gene Ontology analysis of the single nearest genes to (d) bQTLs, (e) caQTLs and (f) clQTLs within 50kb. (g) The Venn diagram shows overlap between bQTL, caQTL and clQTL associated genes.

Further, we used LD R^2^>0.8 instead of exact matching to determine overlap. As expected, the overlaps among bQTLs, caQTLs and clQTLs, and their overlaps with eQTLs all increased (Fig. 4b). To further characterize the associations between eQTLs, bQTLs, caQTLs and clQTLs, we calculated the R^2^ of all the LD pairs within 100 kb maximum distance, and found that the R^2^ distributions of eQTL to significant (*P*<10^−4^) bQTLs, caQTLs or clQTLs was statistically higher than those to non-significant (*P*>0.5) QTLs (Fig. 4c, Additional file 1: Suppl Figs. 4d, 4e and 4f). These data suggest that these QTLs may share a group of the same target genes with eQTLs and could potentially provide a mechanism for their transcriptional regulation.

We next assigned the QTLs to their single nearest genes within 50 kb maximum distance, as the potential target genes. Gene Ontology analysis of the identified genes showed a strong association of these genes with SMC functions, including “extracellular matrix organization”, “blood vessel morphogenesis”, “focal adhesion”, “angiogenesis”, “cell differentiation”, “cell division”, “coronary artery disease”, and “myocardial infarction” (Figs. 4d, 4e and 4f). The targets of these three QTL groups share 98 genes, which have similar gene ontology results as the individual analyses (Fig. 4g, Additional file 1: Suppl Fig. 4g). These data provide a group of novel candidate loci that may contribute to CAD and were examined further in the following studies.

### QTLs located in CAD GWAS causal loci show allele-specific TCF21 binding, chromatin accessibility and chromosomal looping

Given the association of the three types of QTLs with GWAS CAD loci and the QTL target genes with SMC functions, we intersected these CAD SNPs with the QTL colocalized loci and then searched their nearest genes. We found 86 QTLs that associate with 151 CAD SNPs. These SNPs are located in 62 loci across the genome (Additional file 1: Suppl Fig. 5a left). After evaluating the corresponding genes according to published studies, we selected 36 candidate causal genes that are highly likely to be associated with CAD risk. We then separated them into three groups by their distances to CAD SNPs and QTLs (Additional file 1: Suppl Fig. 5a right). Overall, most of these gene were expressed in both GTEx coronary artery tissues (Additional file 1: Suppl Fig. 5b) and HCASMC (Additional file 1: Suppl Fig. 5c). We further investigated the genes in the first group to derive our validation candidates.

These genes, *ARNTL, CCBE1, CDH13*, *EMP1* and *FN1* have been reported as CAD-associated, or related to vascular development or function [2, 18, 28]. In addition, they are involved in epithelial–mesenchymal transition (EMT) processes, which are prominent in vascular disease [29–32]. We validated the QTLs in *CDH13* and *FN1* loci with multiple approaches. For *CDH13,* bQTL rs7198036, and the combination caQTL and clQTL rs12444113 were noted to be located ~10kb downstream of the transcription start site (TSS) (Fig. 5a). We performed ATAC-qPCR and ChIP-qPCR in HCASMC heterozygous lines for these QTLs using Taqman genotyping primers, and confirmed that the G allele had higher chromatin accessibility than the C allele at rs12444113 (Fig. 5b) and allele A had a higher TCF21 occupancy than allele G at rs7198036 (Fig. 5c). To verify the clQTL allele specificity, we employed dCas9-KRAB inhibition “CRISPRi” and investigated *CDH13* expression. The two gRNAs targeting rs12444113 efficiently reduced *CDH13* transcription (Fig. 5d) and chromatin accessibility at the *CDH13* TSS region (Fig. 5e). Since the QTLs are located more than 10kb from the TSS, it is more likely to be a long-range regulatory effect rather than a local *cis*-effect. To further confirm this, we designed an allele-specific 3C-PCR assay to detect the chromosomal interactions with different alleles using HCASMC heterozygous lines. The data confirms that rs12444113 has much greater contact with the *CDH13* TSS than the transcription end site (TES), while they both show allele specificity (Fig. 5f). The measured imbalance for these alleles was consistent with the pre-post allele frequencies in the QTL regression data.

**Figure 5.**
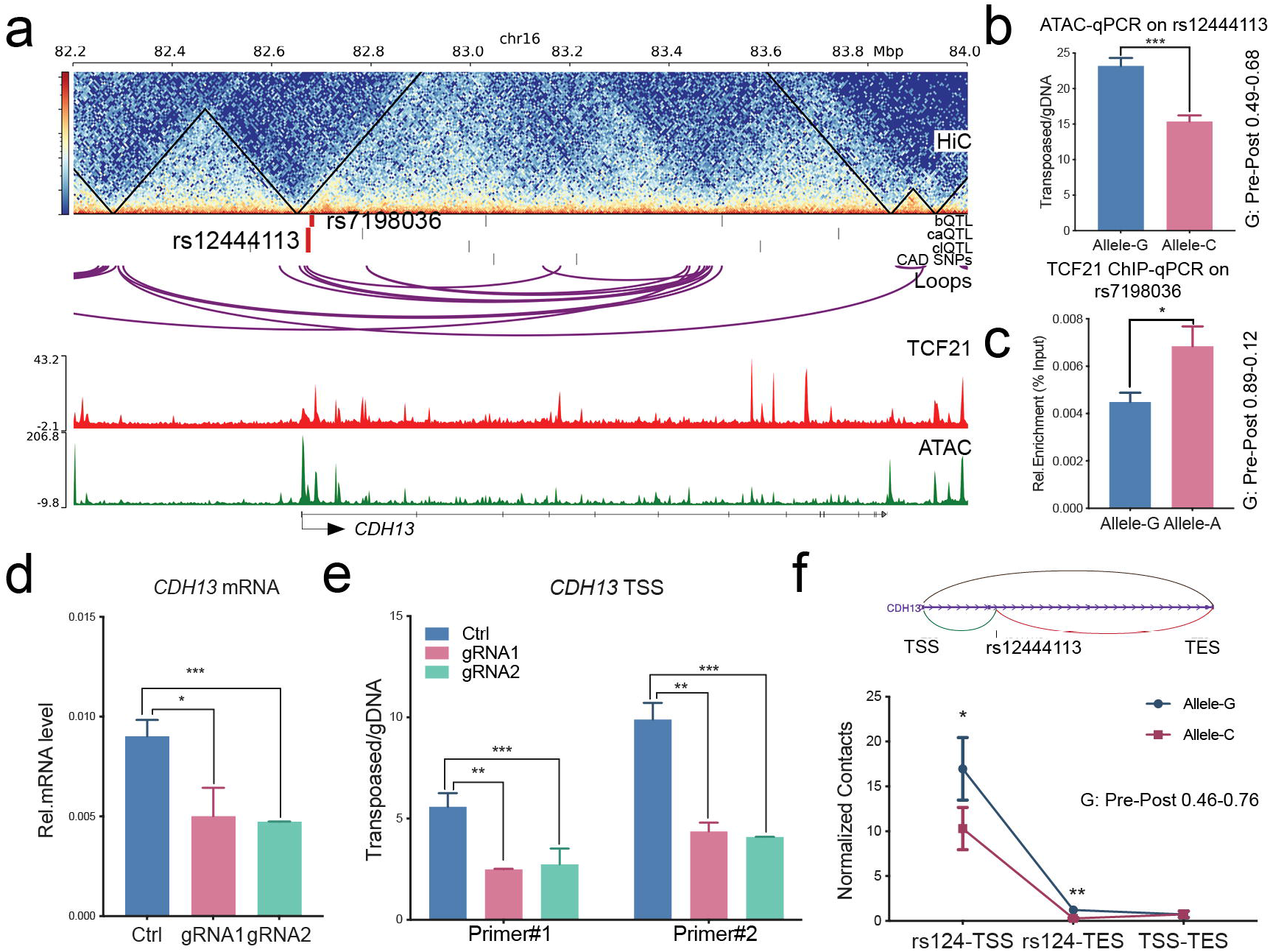
QTLs located in GWAS gene CDH13 show allele-specific TCF21 binding, chromatin accessibility and chromosomal looping. (a) *CDH13* locus, showing bQTL rs7198036 and ca/clQTL rs12444113. Bar graphs show allelic enrichments of (b) chromatin accessibility at rs12444113 and (c) TCF21 binding at rs7198036 identified by allele-specific qPCR, (d) mRNA levels and (e) chromatin accessibility at the TSS region of *CDH13* suppressed by CRISPRi-KRAB targeting at rs12444113. (f) Diagram (top) shows the chromosomal loops between rs12444113 and TSS/TES of *CDH13*. 3C-PCR (bottom) shows the differential chromosomal contacts from rs12444113 to TSS and TES of *CDH13*. Pre-post: allele frequencies in the QTL regression data. Shown are means ± SD; n=3; **** *P*<0.0001; *** *P* <0.001; ** *P* <0.01; * *P* <0.05.

With a similar approach, we validated the caQTL and clQTL rs2692224 variant located ~40kb downstream of the *FN1* TSS (Fig. 6a). Combined ATAC-qPCR (Fig. 6b), CRISPRi (Figs. 6c, 6d) and 3C-PCR (Fig. 6e) data demonstrated that rs2692224 contacts the *FN1* TSS region and allele C has higher chromatin accessibility and chromosomal interaction. Moreover, we further verified the clQTL rs546512774, bQTL and caQTL rs1037169 located in *ARNTL* locus (Additional file 1: Suppl Figs. 6a-6d), clQTL rs1291356, bQTL and caQTL rs7979663 located in *EMP1* locus (Additional file 1: Suppl Figs. 6e-6i), and caQTL, clQTL rs993767 and rs3114275 located in *CCBE1* locus (Additional file 1: Suppl Figs. 6j-6l). All of these QTLs were validated with expected directionalities that are consistent with the pre-post frequencies in QTL regression results.

**Figure 6.**
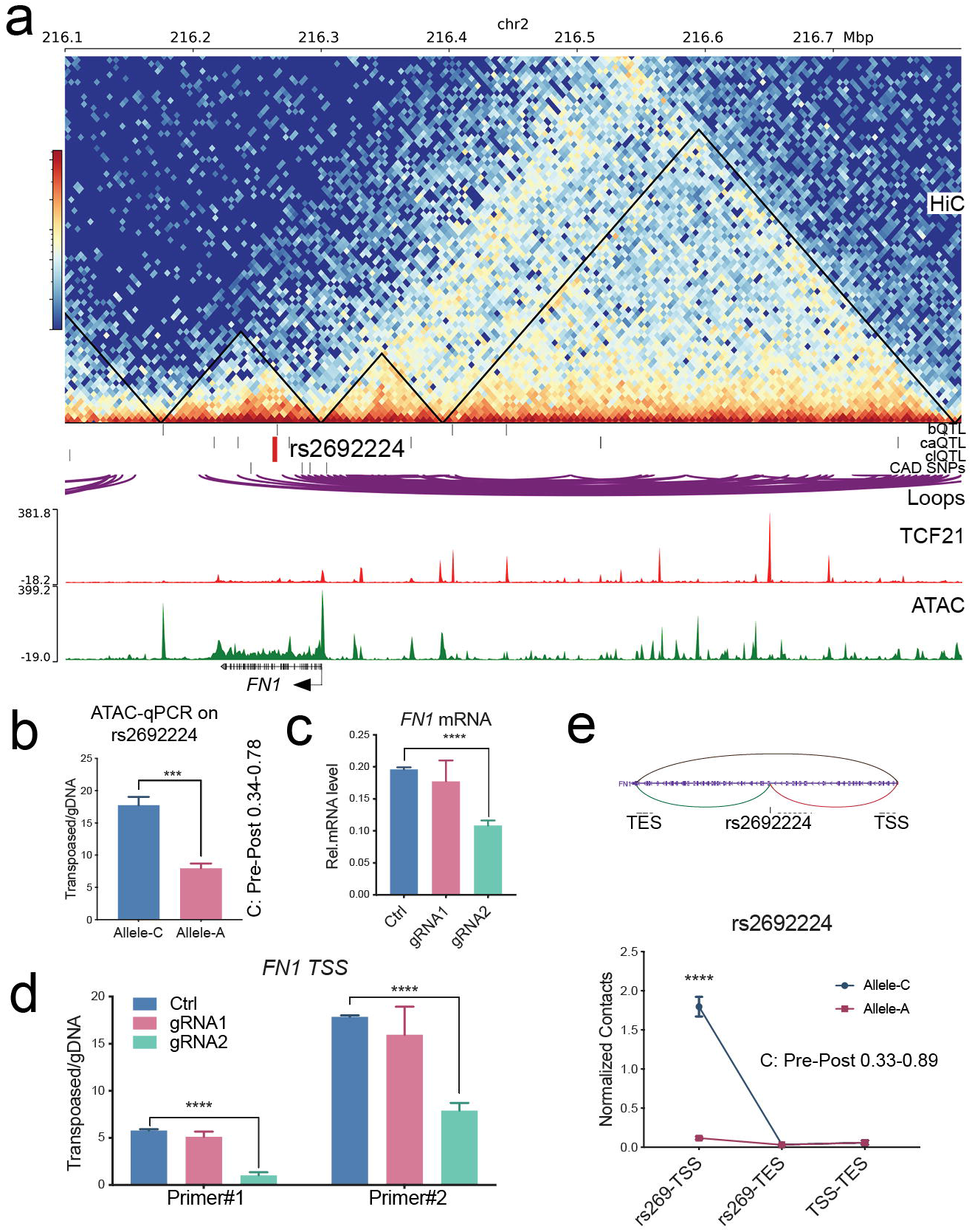
caQTL and clQTL rs2692224 regulates chromatin accessibility and chromosomal interaction at CAD gene FN1. (a) *FN1* locus, showing caQTL and clQTL rs2692224. (b) Bar graphs show allele-specific enrichment of chromatin accessibility at rs2692224 as identified by allele-specific qPCR. (c) mRNA levels and (d) chromatin accessibility at the TSS region of *FN1* were suppressed by CRISPRi-KRAB targeting at rs2692224. (e) Diagram (top) shows the chromosomal loops between rs2692224 and TSS/TES of *FN1*. 3C-PCR (bottom) indicates the differential chromosomal contacts from rs2692224 to TSS of *FN1*. Pre-post: allele frequencies in the QTL regression data. Shown are means ± SD; n=3; **** *P* <0.0001; *** *P* <0.001; ** *P* <0.01; * *P*<0.05.

### Allele-specific looping at the CAD gene *FN1* is regulated by TGFβ1 via response clQTL rs2692224

Fibronectin 1 (FN1) binds to multiple extracellular matrix proteins such as integrin, collagen and fibrin [33]. It regulates cardiovascular development and responds to TGFβ [27, 34, 35]. Given that TGFβ1 and its primary nuclear signaling molecule SMAD3 are both putative CAD-associated genes [23], we sought to investigate a causal mechanism by which this disease related signaling pathway could be linked to *FN1* disease association, through the function of QTLs identified in these studies. As described above, inhibiting the region surrounding rs2692224 with CRISPRi leads to the repression of *FN1* expression. We used TGFβ1 to stimulate multiple HCASMC lines which have different genotypes for rs2692224, including three lines with open allele homozygous C|C, three lines with closed allele homozygous A|A, and two heterozygous A|C lines. We first evaluated *FN1* mRNA levels and found that the increase in expression with TGFβ1 stimulation in the cell lines with the C allele was significantly higher than those with the A allele (Fig. 7a). Similarly, the chromatin accessibility responses to TGFβ at this locus were greater with the rs2692224 C alleles (Fig. 7b). In addition, the 3C-PCR results showed that chromosomal interactions are more active in the cell lines containing the C allele compared with the lines containing the A allele (Fig. 7c). Overall, the C|C homozygous HCASMC are most sensitive to TGFβ1, A|C heterozygous are less sensitive while A|A homozygous are most insensitive to *FN1* expression, chromatin accessibility and chromosomal interaction. These functional studies provide significant evidence that variant rs2692224 serves to link the transcriptional response of the CAD associated gene *FN1* to stimulation by the disease associated TGFβ1 pathway, through mechanisms that regulate chromatin accessibility and chromosomal interactions.

**Figure 7.**
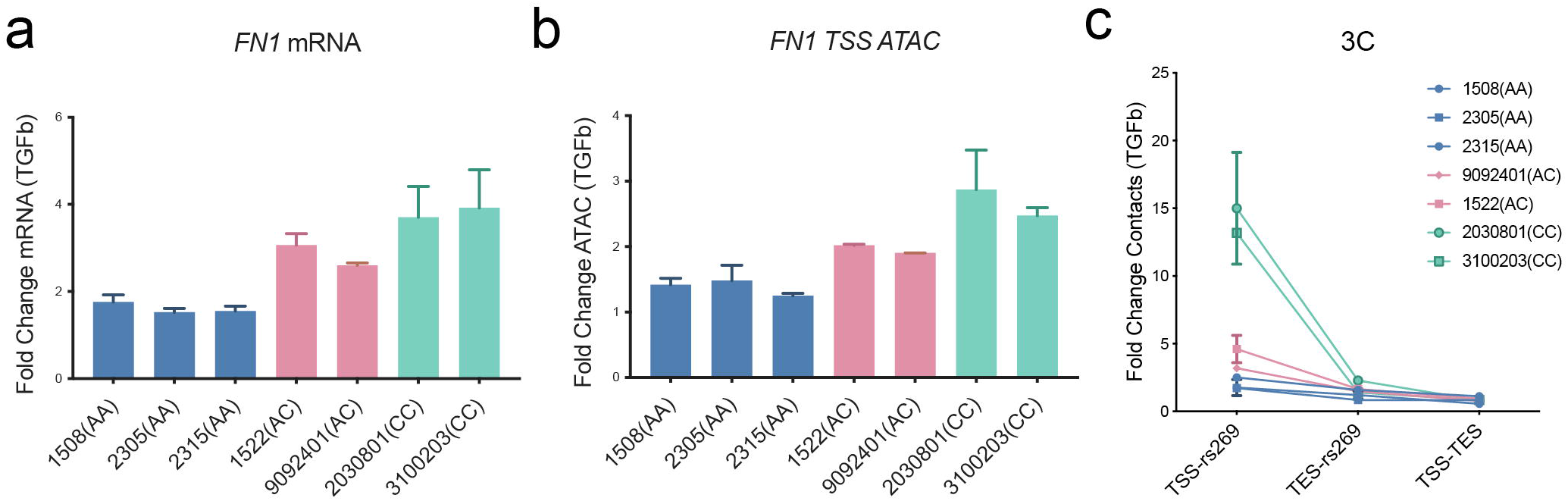
caQTL rs2692224 regulates the allele-specific response of *FN1* to TGFβ signaling. HCASMC lines with different rs2692224 genotypes show distinct activation levels of (a) mRNA expression, (b) chromatin accessibility at the TSS region, and (c) chromosomal contacts between rs2692224 and TSS of *FN1*, in response to TGFβ stimulation. Shown are means ± SD; n=3; **** *P* <0.0001; *** *P* <0.001; ** *P* <0.01; * *P* <0.05.

## Discussion

Recent GWAS have associated thousands of regions of the genome with numerous human traits and complex diseases. For the most part, these associations are likely mediated by variation of gene expression in these loci, due to differences in allelic variation that affects transcription in *cis*. The GWAS data and related insights into the mechanisms of association that are arising from such findings have spurred investigation of the relationships between genetic variation and various genomic features that relate to regulation of transcription [36]. Variants that regulate gene expression, eQTLs, have been the most thoroughly investigated, in cell lines and disease related tissues, and have accelerated the identification of causal genes in associated loci [7, 37–40]. Mapping of QTLs that regulate genomic molecular phenotypes that likely are responsible for eQTL effects on gene expression, such as caQTLs [9] and clQTLs [41], and histone modification, hQTLs [42], have been more limited and restricted primarily to collections of immortalized lymphoblastoid cell lines (LCLs). Studies reported here have significantly expanded our understanding of how these features of the genome regulate gene expression, and how perturbation of DNA sequences that determine these features might contribute to variation in risk for complex human diseases.

Because of our interest in the genetic basis of CAD, and in particular the role of vascular smooth muscle cells in this regard, we have undertaken mapping of genomic regulatory variation in primary cultured HCASMC. Mapping of QTLs in LCLs for instance did not show enrichment in CAD-associated loci, underscoring the importance of studies such as these in disease relevant cell types and tissues [42]. This work was made possible through the pooling approach that dramatically decreases the amount of cell culture work and the expense of multiple assays and sequencing reactions [9, 13]. The individual ChIP-PCR, ATAC-PCR and 3C-PCR assays that we performed represent the most extensive validation of this approach. QTLs of the CAD-associated transcription factor TCF21 were of particular interest because of the enrichment of its binding in other CAD loci [26] and contribution to the risk at these loci where it modulates the epigenome and thus expression of causal genes [15, 27]. We have previously shown that CAD-associated variants are enriched in HCASMC regions of open chromatin [23], and that these cells contribute a significant portion of CAD risk [14]. There was at most 40% overlap of the HCASMC caQTLs with those reported previously for LCLs [9], suggesting that regulatory variation is quite different between these two cell types and underscoring the importance of these smooth muscle cell data for study of vascular disease genetics. We have shown previously that chromosomal looping at CAD loci is regulated by associated allelic variation [16] and mapping of HCASMC clQTLs significantly expands this work with identification of specific variants that are candidate regulators of this genomic feature. To the best of our knowledge this is the first report of mapping clQTLs in primary cultured human cells.

The finding of colocalization of different classes of QTLs in HCASMC is highly informative. It has been established that TF binding can promote open chromatin and create caQTLs [10, 43], and that individual QTLs can influence multiple molecular phenotypes within the same region [42]. In our data there was significant colocalization of bQTLs with caQTLs, and both of these with eQTLs. While the degree of direct overlap of bQTLs and caQTLs was modest (Fig. 4a), there was significant colocalization when variants with LD (R^2^>0.8) were considered (Fig. 4b), and our analyses showed enriched colocalization across the genome for bQTLs and caQTLs (Fig. 4c). Interestingly, both of these classes of variants were also noted to be enriched at JUN binding sites, consistent with a pioneer function for the AP-1 complex (Fig. 3h)[15]. Further, enrichment of these QTLs with the smooth muscle cell MYOCD-SRF complex, SRF binding CArG boxes, indicated a role for these regulatory variants in SMC processes (Fig. 3i). Gene Ontology analysis of genes located nearby to QTLs showed high level enrichment for terms related to SMC processes such as matrix organization, cell-cell interactions - adherence junctions, and vascular development and angiogenesis. By comparison, clQTLs had less colocalization and a different set of enrichment categories, suggesting an independent mechanism from bQTLs and caQTLs.

TF motifs nearby to the bQTLs and clQTLs indicate interactions between the binding and function of these two types of regulatory activities. As expected, there is primarily enrichment for TCF21 and its Class I bHLH dimerization partner TCF12 nearby to the TCF21 bQTLs. Enrichment for AP-1 motifs was identified for both the bQTLs and caQTLs, consistent with its known role in HCASMC [15] and its relationship to lineage determining factors in multiple other cell types [44–46]. TEAD motifs were represented nearby to the bQTL and caQTL regions, and these factors are known to play a significant role in SMC lineage determination and disease risk [47]. Further, caQTLs were associated with SMAD3, a known potent regulator of blood vessel development [48–50]and SMC function [51, 52]as well as CAD association [23, 27]. Interestingly, analysis of the closed caQTLs showed low level enrichment for motifs that mediate binding of ZEB factors, which are known to promote closure of DNA [25].

Perhaps most importantly, mapping these data provide highly useful information for developing credible sets of variants in disease loci implicated by GWAS. For those QTLs that overlap, especially for overlapping bQTLs and caQTLs, there is an increased likelihood that these variants are the active regulatory variant in the region, and thus more likely to contribute to the risk association in the associated locus. However, these QTLs suffer from the same ambiguity regarding functional identity as all variation mapped at a genome-wide level; it is often difficult to discern the functional versus the correlated variants in a haplotype block. In the *CDH13* and *FN1* loci where we have validated the identified QTLs, these QTLs are not in LD with the reported lead SNPs and thus to contribute to disease association would have to be identifying another allele not reported in the GWAS data. The average number of alleles for each GWAS locus is 2-3 and nearby independent alleles are often not reported, so this is a distinct possibility. To link the *FN1* locus QTLs with CAD, we showed that stimulation of HCASMC by TGFβ, a known causal CAD signaling pathway [23, 27, 53], produces a change in the looping pattern as predicted by the clQTL. These molecular QTLs will provide support to fine mapping efforts aimed at identifying disease causal variation, limiting credible sets of genes identified by other methods, and better understanding of the molecular functions of regulatory variation in the human genome.

## Conclusions

In experiments described here we have used a pooling approach with primary cultured HCASMC to map regulatory variation that mediates binding of the CAD associated transcription factor TCF21 (bQTLs) with ChIPseq studies, mapped variation that regulates chromatin accessibility (caQTLs) with ATACseq studies, and chromosomal looping (clQTLs) with HiC methods. We showed that these QTLs are highly associated with CAD GWAS loci and correlated to lead SNPs in these loci, co-localize with smooth muscle cell transcription factor binding and epigenetic modification, show allele-specific function in CAD GWAS loci, and that these functional mapped variants can serve as response QTLs that link expression of causal CAD genes to epigenetic stimulation. Together, these studies represent the most thorough mapping of multiple QTL types in a highly disease relevant primary cultured cell type, and provide novel insights into their functional overlap and mechanisms that underlie these genomic features and their relationship to disease risk.

## Methods

### Primary cell culture and reagents

Primary human coronary artery smooth muscle cells (HCASMCs) derived from normal human donor hearts were purchased from three different manufacturers, Lonza (CC-2583, n=5), PromoCell (C-12511, n=25), Lifeline Cell Tech (FC-0031, n=4), ATCC (PCS-100-021, n=3) and Cell Applications (350-05a, 35) at passage 2 and were maintained in hEGF, insulin, hFGF-b, and 5% FBS supplemented smooth muscle basal media (Lonza # CC-3182) according to the manufacturer’s instructions. All experiments were performed on HCASMCs between passages 4 and 6. Purified rabbit polyclonal antibody against human TCF21 (HPA013189) was purchased from Sigma. Recombinant human TGFβ (AF-100-21C) was purchased from PeproTech and was used at 25 ng/mL for 12 hours after 24 hours serum starvation of the cells.

### ChIP

Briefly, approximately 2×10^7^ pooled HCASMC cells were fixed with 1 % formaldehyde and quenched by glycine. The cells were washed three times with PBS and then harvested in ChIP lysis buffer (50 mM Tris-HCl, pH 8, 5 mM EDTA, 0.5 % SDS). Crosslinked chromatin was sheared for 3×1 min by sonication (Branson SFX250 Sonifier) before extensive centrifugation. Four volumes of ChIP dilution buffer (20 mM Tris-HCl, pH 8.0, 150 mM NaCl, 2 mM EDTA, 1 % Triton X-100) was added to the supernatant. The resulting lysate was then incubated with Dynabeads™ Protein G (Thermo Scientific, 10009D) and antibodies at 4°C overnight. Beads were washed once with buffer 1 (20 mM Tris pH 8, 2 mM EDTA, 150 mM NaCl, 1% Triton X100, 0.1% SDS), once with buffer 2 (10 mM Tris pH 8, 1 mM EDTA, 500 mM NaCl, 1% Triton X100, 0.1% SDS), once with buffer 3 (10 mM Tris pH 8, 1 mM EDTA, 250 mM LiCl, 1% NP40, 1% sodium deoxycholate monohydrate) and twice with TE buffer. DNA was eluted by ChIP elution buffer (0.1 M NaHCO3, 1 % SDS, 20 μg/ml proteinase K). The elution was incubated at 65°C over-night and DNA was extracted with DNA purification kit (Zymo D4013). Library was sequenced on the Illumina HiSeq X instrument to 400 million 150 bp paired end reads.

### ATAC

Approximately 5×10^4^ pooled fresh HCAMSC cells were collected by centrifugation at 500 rcf and washed twice with cold PBS. Nuclei-enriched fractions were extracted with cold resuspension buffer (0.1% NP-40, 0.1% Tween-20, and 0.01% Digitonin), and wash out with 1 ml of cold resuspension buffer containing 0.1% Tween-20 only. Nuclei pellets were collected by centrifugation and resuspended with transposition reaction buffer containing Tn5 transposases (Illumina Nextera TDE1). Transposition reactions were incubated at 37°C for 30 min, followed by DNA purification using the DNA Clean-up and Concentration kit (Zymo D4013). Libraries were amplified using Nextera barcodes and High-Fidelity polymerase (NEB M0541S) and purified using Agencourt Ampure XP beads (Beckman Coulter A63880) double-size selection (0.5X:0.9X). For qPCR experiments, transposed samples were normalized by genomic DNA which was extracted using Quick-DNA Microprep Kit (Zymo D3020). Library was sequenced on the Illumina HiSeq X instrument to 400 million 150 bp paired end reads.

### Hi-C

The Hi-C protocol was modified from HiChIP with excluding the antibody purification [54]. Briefly, approximately 97.5M HCASMC, 1.5M from 65 different cell lines, were pooled and fixed with 1 % formaldehyde and quenched by glycine. The cells were washed with PBS and then harvested in Hi-C lysis buffer (10 mM Tris-HCl, pH 8, 10 mM NaCl, 0.2 % NP-40). Nuclei pellet was incubated in 0.5% SDS at 62°C for 10 min and then quenched by water and 10% Triton X-100. MboI (NEB, R0147) enzyme digested DNA for 2 hours at 37°C. Biotin labeled dATP (Thermo,19524016) and Klenow (NEB, M0210) were used to fill restriction fragment overhangs. The DNA was re-ligated by T4 DNA ligase (NEB, M0202) at room temperature for 4 hours. Re-ligated nuclei were lysed by 0.5% SDS and sheared for 2 min by sonication before extensive centrifugation. Supernatant was incubated at 65°C over-night with Protease K and the DNA was purified. For libraries construction, set aside DNA by 150 ng to the biotin capture step. Biotin labeled DNA was pre-bound to Streptavidin C-1 beads (Thermo, 65-001) in biotin binding buffer. After Tween buffer and TD buffer washing, Tn5 transposition was performed on beads at 55°C for 10 min. Libraries were generated by PCR amplification with Nextera adapters added after washing the beads. Samples were size selected by PAGE purification (300–700 bp) for effective paired end sequencing and adapter dimer removal. All libraries were sequenced on the Illumina HiSeq 4000 instrument to total 800 million 75 bp paired end reads.

### Genotyping and phasing

HCASMC genomic DNA was isolated using DNeasy Blood & Tissue Kit (QIAGEN 69506) and quantified using NanoDrop 1000 Spectrophotometer (Thermo Fisher). Macrogen performed library preparation using Illumina’s TruSeq DNA PCR-Free Library Preparation Kit and 150 bp paired-end sequencing on Illumina HiSeq 4000 instrument.

Whole-genome sequencing data were processed with the *GATK* best practices pipeline [14]. *cutadapt* trimmed reads were aligned to the hg19 reference genome with *bwa*. Duplicate reads in alignment result were marked with *picard markduplicate*. We performed indel realignment and base recalibration with *GATK*. The *GATK HaplotypeCaller* was used to generate gVCF files, which were fed into *GenotypeGVCFs* for joint genotype calling. We recalibrated variants using the *GATK VariantRecalibrato*r module. We further phased our call set with *Beagle*. We first used the *Beagle conform-gt* module to correct any reference genotypes if they are different from hg19. We then phased and imputed against 1000 Genomes phase 3 version 5a. Variants with imputation allelic r^2^ less than 0.8 *and Hardy-Weinberg Equilibrium P*-value less than 1×10^−6^ were filtered out.

### Pooled sequencing data processing

Quality control of pooled ChIPseq data was performed using *fastqc*, and then low-quality bases and adaptor contamination were trimmed by *cutadapt*. Filtered reads were mapped to hg19 genome using *bwa mem* algorithm. Duplicate reads were marked by *picard markduplicate* module and removed with unmapped reads by *samtools*. *macs2 callpeak* was used for peaks calling and input as control. Similar approaches were used for pooled ATACseq with the following modifications. We used *bowtie2* to align reads to hg19 genome. *bedtools* was used for reads format converting and *awk* was used for Tn5 shifting. *macs2 callpeak* with *--broad* was used for peak calling with lambda background.

Pooled HiC paired-end reads were aligned to the HCASMC phasing data masked hg19 genome using the *HiC-Pro* pipeline. Default settings were used to remove duplicate reads, assign reads to MboI restriction fragments, filter for valid interactions, and generate binned interaction matrices. Aligned reads were assigned to a specific allele on the basis of phasing data. *HiC-Pr*o filtered reads were then processed using the *hicpro2juicebox* and *hicpro2fithic* functions. *FitHiC* and *HiCCUPS* ware used to identify high-confidence loops using default parameters. The HiC matrix file was further converted to h5 format by *HiCExplorer hicConvertFormat* module. Genome bias of the matrix was corrected by *hicCorrectMatrix* and *hicFindTADs* was employed to find transcription activation domain (TAD).

### Mapping and analyzing QTLs

Processed alignments of pooled ChIPseq, ATACseq and HiC reads were further realigned by the *hornet* (https://github.com/TheFraserLab/Hornet) pipeline. HCASMC phasing data were combined with 1000 Genome data of CEU population, filtered by MAF>2.5%. Mapped reads that overlap the combined SNPs were identified. For each read that overlaps a SNP, its genotype was swapped with that of the other allele and the read was re-mapped. Re-mapped reads that fail to map to exactly the same location in the genome were discarded.

Pre- and post-allele frequencies, and the resulting p-values, were calculated using published pipeline *cisVar* (https://github.com/TheFraserLab/cisVar). This regression-based approach uses post-allele frequencies together with genotypes of each sample to infer the proportion of each sample in the pool [9, 13]. These proportions will be weighted by any genome-wide differences, since these will be naturally incorporated into the post-frequencies used as input to the regression. In this way, pre-allele frequencies already account for some types of trans-acting variation, increasing the power for mapping *cis*-acting differences.

### GWAS overlap

HCAMSC eQTL data came from a genome-wide association of gene expression with imputed common variation identified in 52 HCASMC lines [14]. CARDIoGRAMplusC4D variants data was from 1000 Genomes-based GWAS meta-analysis [18]. Direct overlap of QTLs and GWAS Catalog SNPs was performed with *bed2GwasCatalogBinomialMod1Ggplot* script from *gwasanalytics* package. The calculation criteria of this script were described previously [55]. We used *LDDirection* (https://github.com/MikeDacre/LDDirection), which employs *plink* and 1000 Genome phasing data of CEU population, to calculate the linkage disequilibrium (LD) between QTL and GWAS SNP pairs within 100kb maximum distance.

### Motif analysis

We used *HOMER findMotifsGenome.pl* script to search for known motifs and to generate *de novo* motifs among the significant QTLs compared to the non-significant QTLs in ±50bp windows. These results were further validated by *MEME*. We also searched the motifs that were differentially enriched among the high-chromatin accessibility or TCF21 binding (open) alleles compared to the low-chromatin accessibility or TCF21 binding (closed) alleles in the shared QTLs. The open allele plus 20 bp on each side (40 bp total) was used as input to *HOMER findMotifs.pl* script, with the closed alleles of the same QTLs used as the background comparison set, or on the opposite direction. Therefore, all significant enrichments are due to QTL variants within the motifs themselves (since any motifs flanking the QTLs would be present in both comparison sets). *PWMScan* was used for position weight matrix scan. We obtained AHR:ARNT (MA0006.1) motif from JASPAR. The JUN and SRF peaks came from the published data [23, 56].

### Cis-regulatory functional enrichment and network analysis

We utilized the *Genomic Regions Enrichment of Annotations Tool (GREAT 3.0)* to search the QTL nearest genes, with the parameter “Single nearest gene”, which was limited to 50kb. Gene ontology analysis from *GREAT* output was done with PANTHER database. Pathways, biological processes, cellular component, phenotype and GAD disease enrichment were carried out using default settings.

### Data visualization

ChIPseq and ATACseq bigWig tracks were converted from filtered alignments using *bedtools* and *UCSC utilities*. HiC matrix generated by *hicpro2juicebox* was visualized by *Juciebox*. The *HiCCUPS* and *FitHiC* output were converted to bigInteract format by custom *awk* command lines. ChIA-PET data came from ENCODE database (ENCSR000CAD and ENCSR361AYD). Genome browser sessions were generated by *WashU Epigenome Browser* or *pyGenomeTracks*.

### CRISPRi validation

We used dCas9 fused to KRAB to knockdown the enhancer region where candidate SNPs were located. The gRNAs targeting SNPs within the maximum 100 bp distance were designed using *Benching* online tools. Synthesized oligos were then cloned into pLV lentivirus vector containing dCas9-KRAB. For virus production, 8.5×10^5^ HEK293T cells were plated in 6-well plate per well. The following day, plasmid encoding lentivirus was co-transfected with pMD2.G and pCMV-dR8.91 into the cells using Lipofectamine 3000 (Thermo Fisher, L3000015) according to the manufacturer’s instructions. ViralBoost Reagent (AlStem Cell Advancements, VB100) was added (1:500) when fresh media was added. Supernatant containing viral particles was collected 72 hours after transfection and filtered. To knockdown, we plated 5×10^4^ HACSMC cells in 6-well plate per well and cultured for 14 hours and then added the virus supernatant to HCASMC for 12 hours with 8 μg/mL polybrene. The cells were cultured for an additional 48 hours for further experiments with media change.

### RNA isolation

RNA for all samples was extracted using the RNeasy mini kit (Qiagen 74106). HCASMC RNA (500 ng) were reverse transcribed using the High capacity RNA-to-cDNA Synthesis kit (Applied Biosystems 4387406).

### Quantitative and genotyping PCR

The purified cDNA or dsDNA samples were assayed by quantitative PCR with ABI ViiA 7 and Power SYBR Green Master Mix (ABI 4368706) using custom designed primers (Additional file 3: Suppl Table. 2). ChIP samples were normalized by input, ATAC transposed samples were normalized by genomic DNA which was extracted using Quick-DNA Microprep Kit (Zymo D3020), and 3C libraries were normalized by post-ligation whole genome DNA without biotin purification. Heterozygous genotypes at the candidate loci were determined using TaqMan SNP genotyping qPCR assays (Thermo Fisher Scientific C 2110683_10, C 8714882_10, C 2042333_10, C 27515710_10, C 29302709_10, C 31103265_10, C 3202579_10). Assays were repeated at least three times. Data shown were average values ± SD of representative experiments.

### Statistical analysis

All experiments were performed by the investigators blinded to the treatments or conditions during the data collection and analysis, using at least two independent biological replicates and treatments/conditions in technical triplicate. R or GraphPad Prism was used for statistical analysis. For motifs and genes enrichment analyses, we used the *cumulative binomial distribution test*. For comparisons between two groups of equal sample size (and assuming equal variance), an unpaired two-tailed *Student’s t-test* was performed or in cases of unequal sample sizes or variance a *Welch’s unequal variances t-test* was performed. *P* values <0.05 were considered statistically significant. For multiple comparison testing, one-way analysis of variance (*ANOVA*) accompanied by *Tukey’s post hoc test* were used as appropriate. All error bars represent standard error of the mean (SE). Number of stars for the *P*-values in the graphs: **** *P*<0.0001; *** *P* <0.001; ** *P* <0.01; * *P* <0.05.

## Abbreviations

GWAS: genome wide association studies
CAD: coronary artery disease
HCASMC: human coronary artery vascular smooth muscle cells
eQTL: expression quantitative trait locus
acQTL: chromatin accessibility quantitative trait locus
clQTL: chromosomal looping quantitative trait locus
LD: linkage disequilibrium
AP-1: activated protein 1
TF: transcription factor
SNP: single nucleotide polymorphism.

## Declarations

### Ethics approval and consent to participate

HCASMC provided by commercial vendors Lonza, PromoCell and Cell Applications, were derived from deceased individuals. Because the donors were deceased, and only deidentified information was provided to the investigators, the work was not considered human research.

### Consent for publication

All authors have reviewed the manuscript and give their consent to publication.

### Availability of data and materials

The datasets generated and analyzed during the current study are available in the Gene Expression Omnibus (GEO), accession number GSE141752 (https://www.ncbi.nlm.nih.gov/geo/query/acc.cgi?acc=GSE141752).

### Author contributions

Q.Z. and T.N performed the experiments. D.I and P.C. helped in the experiments. Q.Z. and M.D. analyzed the data and contributed analyzing scripts. M.P. and B.L. helped in analyzing the data and contributed analyzing scripts. T.Q., Q.Z., T.N. and H.B.F. conceived and designed the experiments. R.W., P.C. and J.B.K discussed the experiments and manuscript. T.Q. and Q.Z. wrote the paper.

## Acknowledgements

We thank members of the Chang and Greenleaf laboratories for help with HiC experiments and M. R Mumbach for assistance interpreting the HiC data. We also thank members of the Fraser laboratory for helpful discussion, and C. Miller at the University of Virginia for help with eQTL studies.

## Additional files

Additional_File_1: Supplemental_Figures.pdf - Supplemental figures and related figure legends.

Additional_File_2: Supplemental_Table_1.xlsx - Regressions of significant TCF21 binding QTLs, chromatin accessibility QTLs and chromosomal looping QTLs.

Additional_File_3: Supplemental_Table_1.xlsx - Customized primers sequences used for real-time qPCR and gRNA sequences.

## Supplementary Figure Legends

**Suppl Figure 1. Quality control of HiC sequencing and interactions.**

(a) The valid pairs called from aligned reads and their orientation statistics. (b) The percentage of valid pairs in all pairs and their classification.

**Suppl Figure 2. QTLs are highly associated with GWAS CAD lead SNPs.**

(a) bQTL, (b) caQTL and (c) clQTL box plots show the fold enrichment of the ratio of LD R^2^>0.8 pairs to the total number of QTL-GWAS SNP pairs, for GWAS CAD-associated diseases.

**Suppl Figure 3. Motif enrichment analysis at clQTLs.**

(a) HOMER reference motif scanning for ±50bp windows around the clQTLs. (b) MEME motif analysis of clQTLs in ±50bp windows with CenrtiMo algorithm.

**Suppl Figure 4. bQTLs and caQTLs are associated with caQTLs in open chromatin regions.**

(a) Total eQTL-eGene pairs and (b) total unique eQTL numbers in HCASMC, compared with the GTEx data. (c) Venn diagram shows the directionality of bQTL and caQTL overlap. Top left: bQTL reference (REF) as bound allele overlap with caQTL REF open allele. Top right: bQTL alternative (ALT) as bound overlap with caQTL ALT open allele. Bottom left: bQTL REF as bound, caQTL ALT open allele. Bottom right: bQTL ALT as bound overlap caQTL REF as open. Box plots show linkage disequilibrium (LD) R^2^ distributions of (d) clQTL, (e) caQTL and (f) bQTL to all other QTL pairs, with non-significant QTLs used as control. (g) Bar graphs show Gene Ontology analysis of the overlapped target genes between bQTL, caQTL and clQTL.

**Suppl Figure 5. Evaluation of CAD causal genes located in bQTL, caQTL and clQTL loci.**

(a) Screening of CAD causal genes located at directly overlapping bQTL, caQTL and clQTL loci. (b) GTEx and c) RNAseq expression in HCASMC lines showing the expression levels of these CAD causal genes.

**Suppl Figure 6. QTLs located in multiple GWAS CAD loci show allele specific TCF21 binding, chromatin accessibility and chromosomal looping.**

(a) ARNTL-TEAD1 locus, showing bQTL rs1037169 and ca/clQTL rs546512774. Bar graphs indicate allelic enrichments of (b) chromatin accessibility and (c) TCF21 binding at rs1037169 identified by allele-specific qPCR. (d) Diagram (top) shows the chromosomal loops between rs546512774 and TSS/TES of ARNTL and TEAD1. 3C-PCR (bottom) showing the differential chromosomal contacts from rs546512774 to TSS and TES of ARNTL. (e) EMP1 locus, showing b/caQTL rs7979663 and clQTL rs1291356. Bar graphs show allelic enrichments of (f) chromatin accessibility, (g) TCF21 binding on rs7979663 and (h) chromosomal contacts at rs1291356 identified by allele-specific qPCR. (i) Diagram (top) shows the chromosomal loops between rs1291356 and TSS/TES of EMP1. 3C-PCR (bottom) showing the differential chromosomal contacts from rs1291356 to TSS and TES of EMP1. (j) CCBE1 locus, showing b/caQTL rs993767 and rs3114275. Bar graphs show allelic enrichments for (k) chromatin accessibility and (l) TCF21 binding on rs993767 and rs3114275 identified by allele-specific qPCR. Pre-post: allele frequencies in the QTL regression data. Shown are means ± SD; n=3; **** P <0.0001; *** P <0.001; ** P <0.01; * P <0.05.

## References

1. Klarin D, Zhu QM, Emdin CA, Chaffin M, Horner S, McMillan BJ, Leed A, Weale ME, Spencer CCA, Aguet F, et al: Genetic analysis in UK Biobank links insulin resistance and transendothelial migration pathways to coronary artery disease. Nat Genet 2017, 49:1392–1397.

2. Nelson CP, Goel A, Butterworth AS, Kanoni S, Webb TR, Marouli E, Zeng L, Ntalla I, Lai FY, Hopewell JC, et al: Association analyses based on false discovery rate implicate new loci for coronary artery disease. Nat Genet 2017, 49:1385–1391.

3. van der Harst P, Verweij N: The Identification of 64 Novel Genetic Loci Provides an Expanded View on the Genetic Architecture of Coronary Artery Disease. Circ Res 2017, 122:433–443.

4. Battle A, Mostafavi S, Zhu X, Potash JB, Weissman MM, McCormick C, Haudenschild CD, Beckman KB, Shi J, Mei R, et al: Characterizing the genetic basis of transcriptome diversity through RNA-sequencing of 922 individuals. Genome Res 2013.

5. Conde L, Bracci PM, Richardson R, Montgomery SB, Skibola CF: Integrating GWAS and Expression Data for Functional Characterization of Disease-Associated SNPs: An Application to Follicular Lymphoma. Am J Hum Genet 2013, 92:126–130.

6. Consortium GT, Laboratory DA, Coordinating Center-Analysis Working G, Statistical Methods groups-Analysis Working G, Enhancing Gg, Fund NIHC, Nih/Nci, Nih/Nhgri, Nih/Nimh, Nih/Nida, et al: Genetic effects on gene expression across human tissues. Nature 2017, 550:204–213.

7. Lappalainen T, Sammeth M, Friedlander MR, t Hoen PA, Monlong J, Rivas MA, Gonzalez-Porta M, Kurbatova N, Griebel T, Ferreira PG, et al: Transcriptome and genome sequencing uncovers functional variation in humans. Nature 2013, 501:506–511.

8. Maurano MT, Humbert R, Rynes E, Thurman RE, Haugen E, Wang H, Reynolds AP, Sandstrom R, Qu H, Brody J, et al: Systematic localization of common disease-associated variation in regulatory DNA. Science 2012, 337:1190–1195.

9. Tehranchi A, Hie B, Dacre M, Kaplow I, Pettie K, Combs P, Fraser HB: Fine-mapping cis-regulatory variants in diverse human populations. Elife 2019, 8.

10. Degner JF, Pai AA, Pique-Regi R, Veyrieras JB, Gaffney DJ, Pickrell JK, De Leon S, Michelini K, Lewellen N, Crawford GE, et al: DNase I sensitivity QTLs are a major determinant of human expression variation. Nature 2012, 482:390–394.

11. Greenwald WW, Li H, Benaglio P, Jakubosky D, Matsui H, Schmitt A, Selvaraj S, D’Antonio M, D’Antonio-Chronowska A, Smith EN, Frazer KA: Subtle changes in chromatin loop contact propensity are associated with differential gene regulation and expression. Nat Commun 2019, 10:1054.

12. Gorkin DU, Qiu Y, Hu M, Fletez-Brant K, Liu T, Schmitt AD, Noor A, Chiou J, Gaulton KJ, Sebat J, et al: Common DNA sequence variation influences 3-dimensional conformation of the human genome. Genome Biol 2019, 20:255.

13. Tehranchi AK, Myrthil M, Martin T, Hie BL, Golan D, Fraser HB: Pooled ChIP-Seq Links Variation in Transcription Factor Binding to Complex Disease Risk. Cell 2016, 165:730–741.

14. Liu B, Pjanic M, Wang T, Nguyen T, Gloudemans M, Rao A, Castano VG, Nurnberg S, Rader DJ, Elwyn S, et al: Genetic Regulatory Mechanisms of Smooth Muscle Cells Map to Coronary Artery Disease Risk Loci. Am J Hum Genet 2018, 103:377–388.

15. Zhao Q, Wirka R, Nguyen T, Nagao M, Cheng P, Miller CL, Kim JB, Pjanic M, Quertermous T: TCF21 and AP-1 interact through epigenetic modifications to regulate coronary artery disease gene expression. Genome Med 2019, 11:23.

16. Mumbach MR, Satpathy AT, Boyle EA, Dai C, Gowen BG, Cho SW, Nguyen ML, Rubin AJ, Granja JM, Kazane KR, et al: Enhancer connectome in primary human cells identifies target genes of disease-associated DNA elements. Nat Genet 2017, 49:1602–1612.

17. Buniello A, MacArthur JAL, Cerezo M, Harris LW, Hayhurst J, Malangone C, McMahon A, Morales J, Mountjoy E, Sollis E, et al: The NHGRI-EBI GWAS Catalog of published genome-wide association studies, targeted arrays and summary statistics 2019. Nucleic Acids Res 2019, 47:D1005–D1012.

18. Nikpay M, Goel A, Won HH, Hall LM, Willenborg C, Kanoni S, Saleheen D, Kyriakou T, Nelson CP, Hopewell JC, et al: A comprehensive 1,000 Genomes-based genome-wide association meta-analysis of coronary artery disease. Nat Genet 2015, 47:1121–1130.

19. Liang J, Le TH, Edwards DRV, Tayo BO, Gaulton KJ, Smith JA, Lu Y, Jensen RA, Chen G, Yanek LR, et al: Single-trait and multi-trait genome-wide association analyses identify novel loci for blood pressure in African-ancestry populations. PLoS Genet 2017, 13:e1006728.

20. Fujimaki T, Oguri M, Horibe H, Kato K, Matsuoka R, Abe S, Tokoro F, Arai M, Noda T, Watanabe S, Yamada Y: Association of a transcription factor 21 gene polymorphism with hypertension. Biomed Rep 2015, 3:118–122.

21. Wang Y, Wang L, Liu X, Zhang Y, Yu L, Zhang F, Liu L, Cai J, Yang X, Wang X: Genetic variants associated with myocardial infarction and the risk factors in Chinese population. PLoS One 2014, 9:e86332.

22. Wang J, Gao X, Wang M, Zhang J: Clinicopathological significance and biological role of TCF21 mRNA in breast cancer. Tumour Biol 2015, 36:8679–8683.

23. Miller CL, Pjanic M, Wang T, Nguyen T, Cohain A, Perisic L, Hedin U, Betsholtz C, Ruusalepp A, Franzen O, et al: Integrative functional genomics identifies regulatory mechanisms at coronary artery disease loci. Nat Commun 2016:12092–12108.

24. Khan A, Fornes O, Stigliani A, Gheorghe M, Castro-Mondragon JA, van der Lee R, Bessy A, Cheneby J, Kulkarni SR, Tan G, et al: JASPAR 2018: update of the open-access database of transcription factor binding profiles and its web framework. Nucleic Acids Res 2018, 46:D260–D266.

25. Wu LM, Wang J, Conidi A, Zhao C, Wang H, Ford Z, Zhang L, Zweier C, Ayee BG, Maurel P, et al: Zeb2 recruits HDAC-NuRD to inhibit Notch and controls Schwann cell differentiation and remyelination. Nat Neurosci 2016, 19:1060–1072.

26. Sazonova O, Zhao Y, Nurnberg S, Miller C, Pjanic M, Castano VG, Kim JB, Salfati EL, Kundaje AB, Bejerano G, et al: Characterization of TCF21 downstream target regions identifies a transcriptional network linking multiple independent coronary artery disease loci. PLoS Genet 2015, 11.

27. Iyer D, Zhao Q, Wirka R, Naravane A, Nguyen T, Liu B, Nagao M, Cheng P, Miller CL, Kim JB, et al: Coronary artery disease genes SMAD3 and TCF21 promote opposing interactive genetic programs that regulate smooth muscle cell differentiation and disease risk. PLoS Genet 2018, 14:e1007681.

28. Bonet F, Pereira PNG, Bover O, Marques S, Inacio JM, Belo JA: CCBE1 is required for coronary vessel development and proper coronary artery stem formation in the mouse heart. Dev Dyn 2018, 247:1135–1145.

29. Mesci A, Huang X, Taeb S, Jahangiri S, Kim Y, Fokas E, Bruce J, Leong HS, Liu SK: Targeting of CCBE1 by miR-330-3p in human breast cancer promotes metastasis. Br J Cancer 2017, 116:1350–1357.

30. Li CL, Yang D, Cao X, Wang F, Hong DY, Wang J, Shen XC, Chen Y: Fibronectin induces epithelial-mesenchymal transition in human breast cancer MCF-7 cells via activation of calpain. Oncol Lett 2017, 13:3889–3895.

31. Xu X, Zhou Y, Xie C, Wei SM, Gan H, He S, Wang F, Xu L, Lu J, Dai W, et al: Genome-wide screening reveals an EMT molecular network mediated by Sonic hedgehog-Gli1 signaling in pancreatic cancer cells. PLoS One 2012, 7:e43119.

32. Tahara T, Shibata T, Okubo M, Ishizuka T, Nakamura M, Nagasaka M, Nakagawa Y, Ohmiya N, Arisawa T, Hirata I: DNA methylation status of epithelial-mesenchymal transition (EMT)--related genes is associated with severe clinical phenotypes in ulcerative colitis (UC). PLoS One 2014, 9:e107947.

33. Pankov R, Yamada KM: Fibronectin at a glance. J Cell Sci 2002, 115:3861–3863.

34. Chen D, Wang X, Liang D, Gordon J, Mittal A, Manley N, Degenhardt K, Astrof S: Fibronectin signals through integrin alpha5beta1 to regulate cardiovascular development in a cell type-specific manner. Dev Biol 2015, 407:195–210.

35. Keski-Oja J, Raghow R, Sawdey M, Loskutoff DJ, Postlethwaite AE, Kang AH, Moses HL: Regulation of mRNAs for type-1 plasminogen activator inhibitor, fibronectin, and type I procollagen by transforming growth factor-beta. Divergent responses in lung fibroblasts and carcinoma cells. J Biol Chem 1988, 263:3111–3115.

36. Neumeyer S, Hemani G, Zeggini E: Strengthening Causal Inference for Complex Disease Using Molecular Quantitative Trait Loci. Trends Mol Med 2019.

37. Zhu Z, Zhang F, Hu H, Bakshi A, Robinson MR, Powell JE, Montgomery GW, Goddard ME, Wray NR, Visscher PM, Yang J: Integration of summary data from GWAS and eQTL studies predicts complex trait gene targets. Nat Genet 2016, 48:481–487.

38. Stranger BE, Nica AC, Forrest MS, Dimas A, Bird CP, Beazley C, Ingle CE, Dunning M, Flicek P, Koller D, et al: Population genomics of human gene expression. Nat Genet 2007, 39:1217–1224.

39. Nica AC, Montgomery SB, Dimas AS, Stranger BE, Beazley C, Barroso I, Dermitzakis ET: Candidate causal regulatory effects by integration of expression QTLs with complex trait genetic associations. PLoS Genet 2010, 6:e1000895.

40. Montgomery SB, Dermitzakis ET: The resolution of the genetics of gene expression. Hum Mol Genet 2009, 18:R211–215.

41. Gorkin DU, Qiu Y, Hu M, Fletez-Brant K, Liu T, Schmitt AD, Noor A, Chiou J, Gaulton KJ, Sebat J, et al: Common DNA sequence variation influences 3-dimensional conformation of the human genome. bioRxiv 2019.

42. Grubert F, Zaugg JB, Kasowski M, Ursu O, Spacek DV, Martin AR, Greenside P, Srivas R, Phanstiel DH, Pekowska A, et al: Genetic Control of Chromatin States in Humans Involves Local and Distal Chromosomal Interactions. Cell 2015, 162:1051–1065.

43. Kumasaka N, Knights AJ, Gaffney DJ: Fine-mapping cellular QTLs with RASQUAL and ATAC-seq. Nat Genet 2015.

44. Heinz S, Benner C, Spann N, Bertolino E, Lin YC, Laslo P, Cheng JX, Murre C, Singh H, Glass CK: Simple combinations of lineage-determining transcription factors prime cis-regulatory elements required for macrophage and B cell identities. Mol Cell 2010, 38:576–589.

45. Heinz S, Romanoski CE, Benner C, Allison KA, Kaikkonen MU, Orozco LD, Glass CK: Effect of natural genetic variation on enhancer selection and function. Nature 2013, 503:487–492.

46. Romanoski CE, Che N, Yin F, Mai N, Pouldar D, Civelek M, Pan C, Lee S, Vakili L, Yang WP, et al: Network for activation of human endothelial cells by oxidized phospholipids: a critical role of heme oxygenase 1. Circ Res 2011, 109:e27–41.

47. Almontashiri NA, Antoine D, Zhou X, Vilmundarson RO, Zhang SX, Hao KN, Chen HH, Stewart AF: 9p21.3 Coronary Artery Disease Risk Variants Disrupt TEAD Transcription Factor-Dependent TGFbeta Regulation of p16 Expression in Human Aortic Smooth Muscle Cells. Circulation 2015.

48. Agah R, Prasad KS, Linnemann R, Firpo MT, Quertermous T, Dichek DA: Cardiovascular overexpression of transforming growth factor-beta(1) causes abnormal yolk sac vasculogenesis and early embryonic death. Circ Res 2000, 86:1024–1030.

49. Gaengel K, Genove G, Armulik A, Betsholtz C: Endothelial-mural cell signaling in vascular development and angiogenesis. Arterioscler Thromb Vasc Biol 2009, 29:630–638.

50. Kurpinski K, Lam H, Chu J, Wang A, Kim A, Tsay E, Agrawal S, Schaffer DV, Li S: Transforming growth factor-beta and notch signaling mediate stem cell differentiation into smooth muscle cells. Stem Cells 2010, 28:734–742.

51. Reddy KB, Howe PH: Transforming growth factor beta 1-mediated inhibition of smooth muscle cell proliferation is associated with a late G1 cell cycle arrest. J Cell Physiol 1993, 156:48–55.

52. Suwanabol PA, Seedial SM, Zhang F, Shi X, Si Y, Liu B, Kent KC: TGF-beta and Smad3 modulate PI3K/Akt signaling pathway in vascular smooth muscle cells. Am J Physiol Heart Circ Physiol 2012, 302:H2211–2219.

53. Zeng L, Dang TA, Schunkert H: Genetics links between transforming growth factor beta pathway and coronary disease. Atherosclerosis 2016, 253:237–246.

54. Mumbach MR, Rubin AJ, Flynn RA, Dai C, Khavari PA, Greenleaf WJ, Chang HY: HiChIP: efficient and sensitive analysis of protein-directed genome architecture. Nat Methods 2016, 13:919–922.

55. Kim BJ, Pjanic M, Nguyen T, Miller CL, Liu B, Wang T, Sazonova O, Carcamo-Orive I, Perisic L, Maegdefessel L, et al: TCF21 and the aryl-hydrocarbon receptor cooperate to activate a pro-atherosclerotic gene expression program. PLoS Genet 2017, 13.:1006750.

56. Nagao M, Zhao Q, Wirka R, Bagga J, Nguyen T, Cheng P, Kim JB, Pjanic M, Miano JM, Quertermous T: Coronary disease associated gene TCF21 inhibits smooth muscle cell differentiation by blocking the myocardin-serum response factor pathway. Circ Res 2019, in press.

