## Supplemental Figures for "Quantitative trait loci mapped for TCF21 binding, chromatin accessibility and chromosomal looping in coronary artery smooth muscle cells reveal molecular mechanisms of coronary disease loci"

Supplemental Figure.1

a

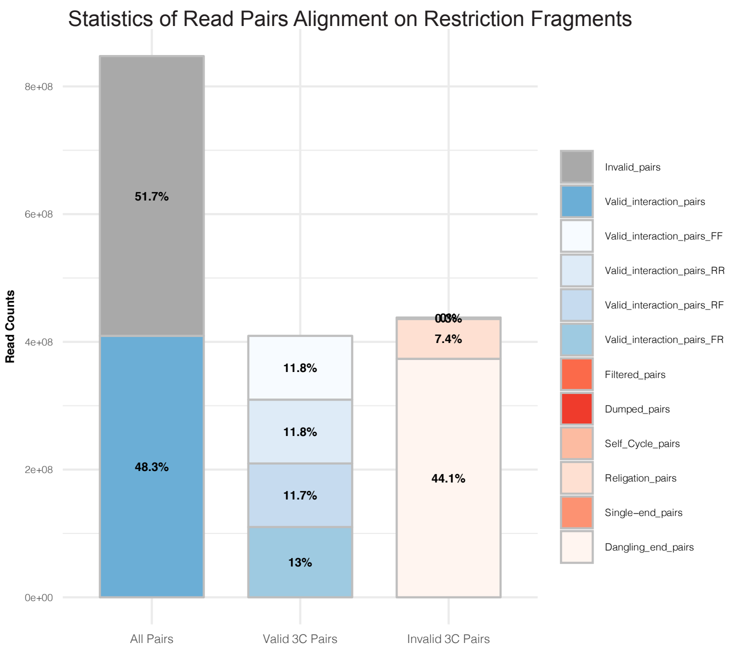


b

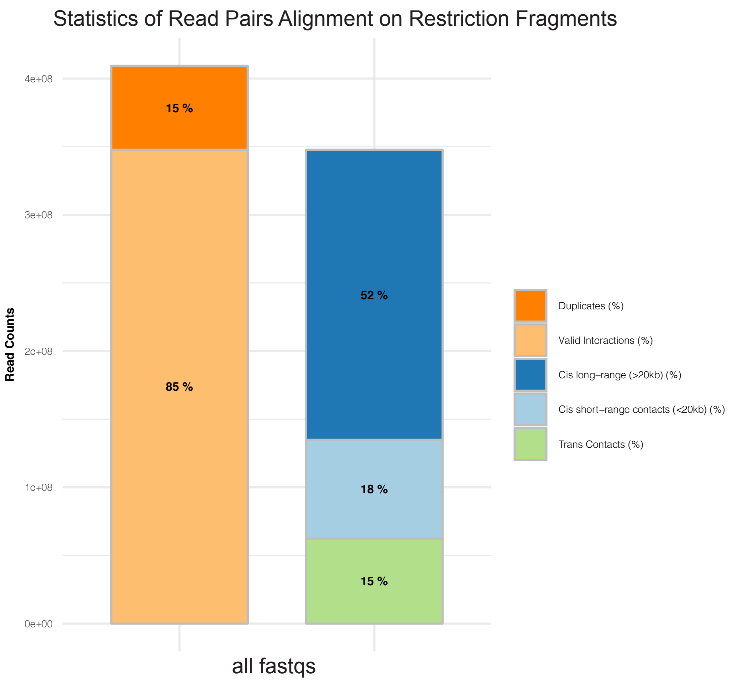

Supplemental Figure.2

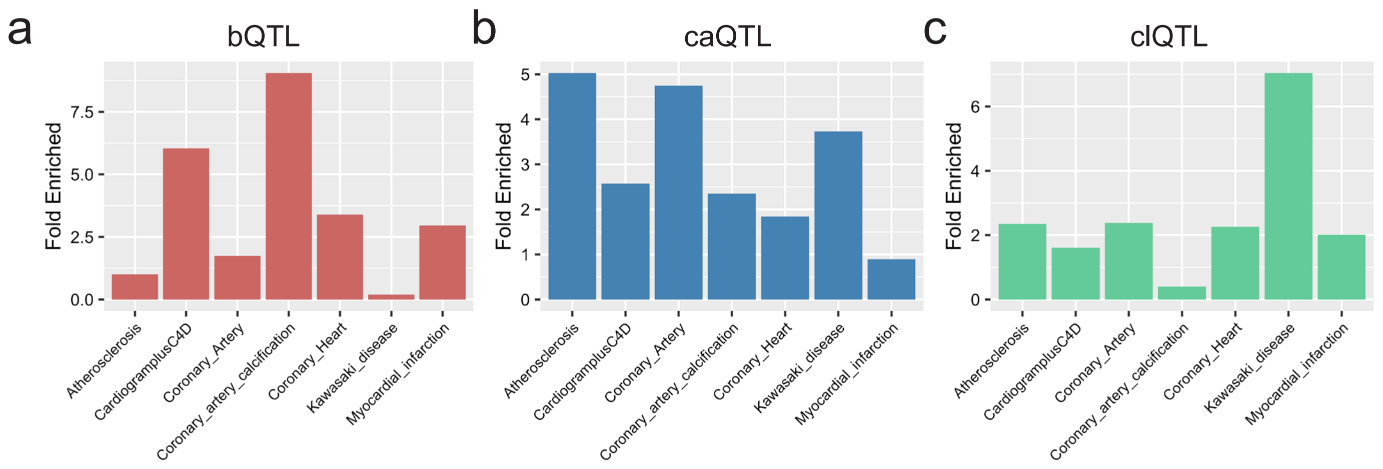

a

cIQTL

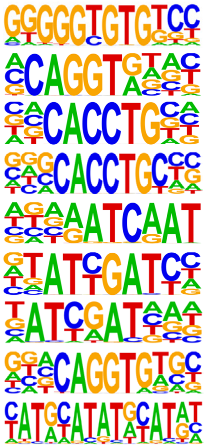

| | Motif | Rank | *P* value |
|---|---|---|---|
| KLF10 | | 1 | 1e-29 |
| ZEB1 | | 2 | 1e-18 |
| E2A | | 3 | 1e-16 |
| Slug | | 4 | 1e-12 |
| Cux2 | | 5 | 1e-11 |
| HNF6 | | 6 | 1e-10 |
| CUX1 | | 7 | 1e-10 |
| ZEB2 | | 8 | 1e-10 |
| OCT | | 9 | 1e-8 |

b

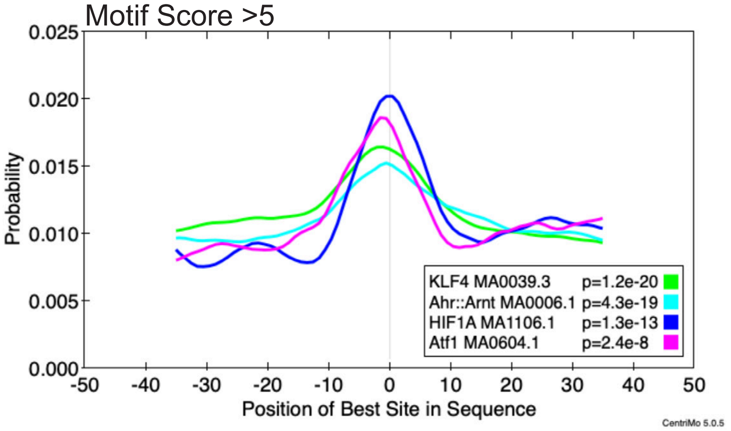

### Supplemental Figure.4

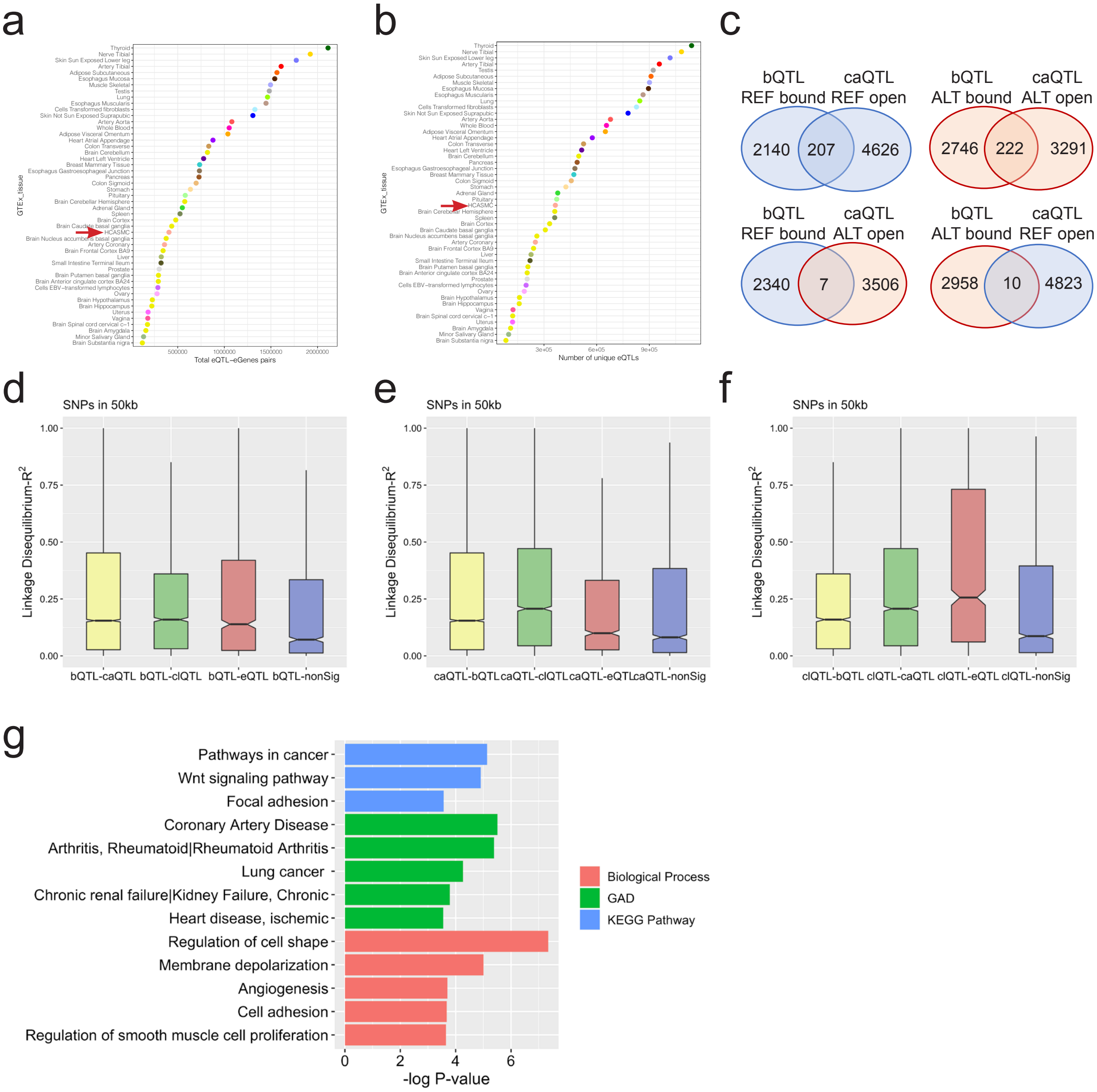

Supplemental Figure.5

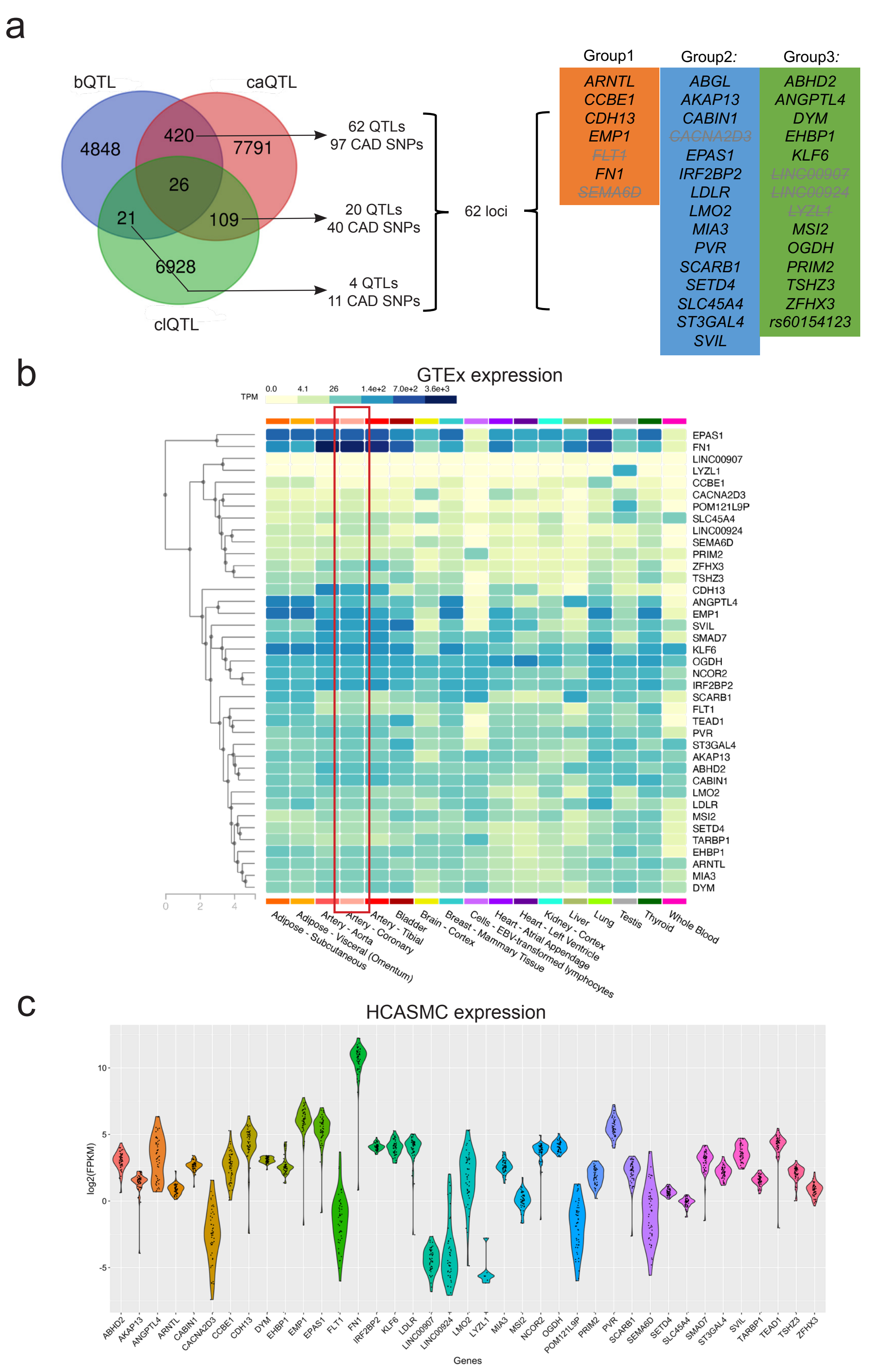

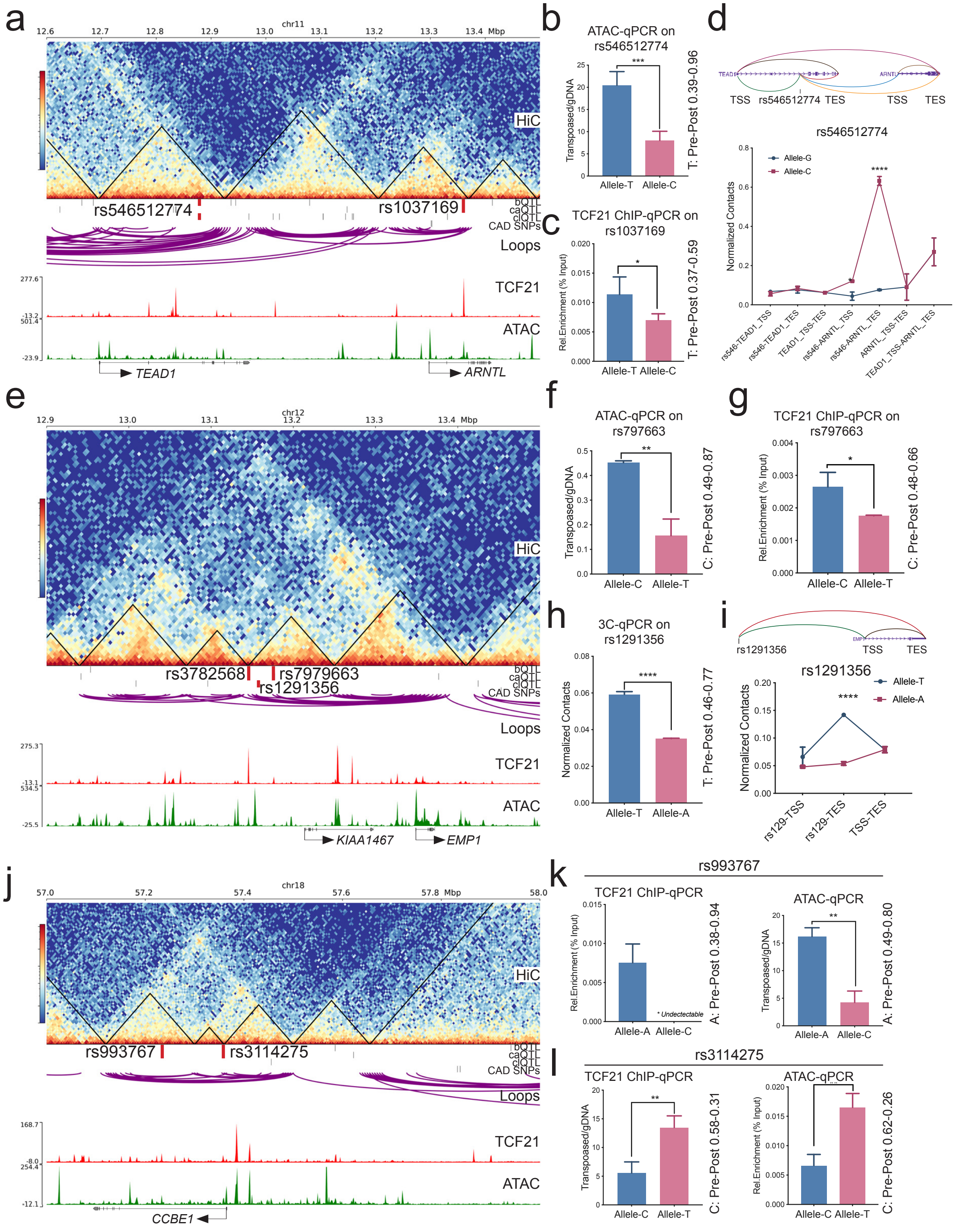
a
b
c
d
e
f
g
h
i
j
k
l
chr11
HiC
rs546512774
rs1037169
bQTL
caQTL
clQTL
CAD SNPs
Loops
TCF21
ATAC
TEAD1
ARNTL
ATAC-qPCR on rs546512774
Transposaed/gDNA
Allele-T
Allele-C
T: Pre-Post 0.39-0.96
TCF21 ChIP-qPCR on rs1037169
Rel.Enrichment (% Input)
T: Pre-Post 0.37-0.59
TSS
rs546512774
TES
Normalized Contacts
Allele-G
Allele-C
rs546-TEAD1_TSS
rs546-TEAD1_TES
TEAD1_TSS-TES
rs546-ARNTL_TSS
rs546-ARNTL_TES
ARNTL_TSS-TES
TEAD1_TSS-ARNTL_TES
chr12
rs3782568
rs7979663
rs1291356
KIAA1467
EMP1
ATAC-qPCR on rs797663
C: Pre-Post 0.49-0.87
TCF21 ChIP-qPCR on rs797663
C: Pre-Post 0.48-0.66
3C-qPCR on rs1291356
Allele-A
T: Pre-Post 0.46-0.77
rs1291356
rs129-TSS
rs129-TES
TSS-TES
chr18
rs993767
rs3114275
CCBE1
rs993767
TCF21 ChIP-qPCR
ATAC-qPCR
* Undectectable
A: Pre-Post 0.38-0.94
A: Pre-Post 0.49-0.80
rs3114275
C: Pre-Post 0.58-0.31
C: Pre-Post 0.62-0.26
